# SETD3 protein is the actin-specific histidine *N*-methyltransferase

**DOI:** 10.1101/266882

**Authors:** Sebastian Kwiatkowski, Agnieszka K. Seliga, Maria Veiga-da-Cunha, Didier Vertommen, Marianna Terreri, Takao Ishikawa, Iwona Grabowska, Adam K Jagielski, Jakub Drozak

**Author notes:** S.K. and A.K.S. contributed equally to this work. To whom correspondence should be addressed: Jakub Drozak, Department of Metabolic Regulation, Institute of Biochemistry, Faculty of Biology, University of Warsaw, Miecznikowa 1, 02-096 Warsaw, Poland.

## Abstract

Protein histidine methylation is rarely studied posttranslational modification of unknown biochemical importance. In vertebrates, only a few methylhistidne-containing proteins have been reported so far, including β-actin as an essential example. The evolutionary conserved methylation of β-actin H73 residue is catalyzed by a specific histidine *N*-methyltransferase that has never been identified molecularly. In the present investigation, we have purified actin-specific histidine *N*-methyltransferase from rat muscles about 1200-fold. Its activity was studied by the radiochemical assay employing either homogeneous recombinant human β-actin produced in *E. coli* or its mutated form exhibiting substitution of H73 by Ala residue (H73A) as substrates. Three polypeptides of ≈65, 75 and 90 kDa coeluting with the enzyme activity were identified in the preparation. Mass spectrometry analysis of these polypeptides resulted in the identification of SETD3 methyltransferase as the only plausible candidate. Rat SETD3 and its human ortholog were expressed in COS-7 cells, purified to homogeneity and shown to catalyze methylation of β-actin at H73 residue as confirmed by mass spectrometry analysis. The SETD3 enzyme was active towards a synthetic peptide corresponding to residues 69-77 of β-actin, but not to its mutated form exhibiting His-to-Ala substitution. Finally, Setd3-deficient HAP1 cells were devoid of methylated H73 in β-actin and exhibited phenotypic changes, including a decrease in F-actin content and an increased glycolytic activity. We conclude that SETD3 is the actin-specific histidine *N*-methyltransferase. The data show for the first time the molecular identity of protein histidine N-methyltransferase in vertebrates and throw new light on the substrate specificity of SET-domain-containing enzymes.

## INTRODUCTION

Protein methylation is one of the most common post-translation modification (PTM) in eukaryotic cells [for review see Clarke, 2013] and histones are probably the best studied substrates of protein *N*-methyltransferases. Much is therefore known about methylation of lysine and arginine residues in histones as well as the fundamental role of such modifications in the epigenetic control of mammalian gene expression [Greer and Shi, 2012]. However, an increasing number of studies show methylation of non-histone proteins as a prevalent PTM that regulates diverse biological processes, including protein synthesis and signal transduction [for review see Biggar and Li, 2015]. More interestingly, non-histone proteins may be modified on atypical sites for methylation such as glutamate, glutamine, cysteine, and histidine residues [Clarke, 2013]. Methylation of histidine residues has been long reported just for a few proteins in Nature, including β-actin [Johnson *et al.*, 1967], S100A9 protein [Raftery *et al.*, 1996], myosin [Elzinga and Collins, 1977] and its kinase [Meyer i Mayr, 1987], and ribosomal protein RPL3 [Webb *et al.*, 2010], whereas a recent study of Ning and coworkers [2016] suggests that such modification may be a quite common phenomenon, involving dozens if not hundreds of intracellular proteins in mammalian cells.

Knowledge of protein histidine *N*-methyltransferases is rather limited, and yeast YIL110W (Hpm1p) protein, which catalyzes methylation of histidine 243 in ribosomal RPL3 protein, remains the first and only histidine protein methyltransferase molecularly identified up to date [Webb *et al*, 2010]. The enzyme responsible for actin histidine methylation has been partially purified, but no gene encoding this enzyme has been identified yet [Vijayasarathy and Rao, 1987; Raghavan *et al.*, 1992].

Actin is one of the most abundant protein in eukaryotic cells and a major component of the cytoskeleton [for review, Pollard and Cooper, 2009]. It is highly evolutionary conserved (≈90% identity between yeast and human protein), and in vertebrates, three main isoforms of this protein differing by only a few amino acids at their N-terminus have been identified. α-Actins are expressed in skeletal, cardiac and smooth muscle, while β- and γ-isoforms are present in nonmuscle and muscle cells. Under physiological conditions, actin exists as a 42-kDa monomeric globular protein (G-actin) that binds ATP and spontaneously polymerize into stable filaments (F-actin). Once filament has been assembled, the terminal phosphate of the bound ATP is hydrolyzed and slowly released from the protein, leading to the sequential depolymerization of the filament. The maintenance of a pool of actin monomers, the initialization of polymerization process as well as the regulation of assembly and turnover of filaments are controlled and tuned by more than 100 interacting partner proteins, making actin to contribute to more protein-protein interactions than any other protein does [Lappalainen, 2016].

As actin is essential for the survival of cells, participating in numerous fundamental cellular processes, including cell growth and division, and cellular motility [Dominguez and Holmes, 2011], an attention has been devoted to elucidate the role of the conserved methylation of H73 residue [Nyman *et al.*, 2002]. It was shown that H73 methylation stabilizes actin filaments, while the absence of such modification results in an increased rate of filaments depolymerization [Nyman *et al.*, 2002]. No physiological effects of actin hypomethylation have yet been identified in cells, tissues or organisms.

In the current investigation, we report the identification of actin-specific histidine *N*-methyltransferase (EC 2.1.1.85) designated SET domain-containing protein 3 (SETD3, C14orf154) that has been previously shown to act as histone lysine methyltransferase [Eom *et al.*, 2011]. We have characterized this enzyme and proved that it specifically modifies H73 residue of human (mammalian) β-actin. We have also shown that a loss of SETD3 activity is accompanied by changes of human HAP1 cells phenotype, including a decrease in F-actin content and an increased glycolytic activity in HAP1 cells. The identification of SETD3 protein as actin-specific histidine *N*-methyltransferase is the first report of enzyme that catalyzes such post-translational modification in vertebrates. This finding throws also a new light on the substrate specificity of SET-domain enzymes, as all these enzymes characterized so far are protein-lysine methyltransferases [Clarke, 2013].

## EXPERIMENTAL PROCEDURES

### Materials

Reagents, of analytical grade whenever possible, were from Sigma-Aldrich (Saint Louis, USA), Roche Diagnostics (Basel, Switzerland.) or Merck (Darmstadt, Germany). S-[methyl-^3^H]adenosyl-L-methionine ([^3^H]SAM) and Ultima Gold LSC cocktail were purchased from PerkinElmer (Waltham, USA), while S-[methyl-^2^H]adenosyl-L-methionine ([^2^H]SAM) came from C/D/N Isotopes Inc. (Pointe-Claire, Canada). Peptides were from Caslo Laboratory (Lyngby, Denmark). DEAE-Sepharose, Q-Sepharose, Phenyl-Sepharose, Superdex 200, HiScreen Blue FF, HisTrap HP (Ni^2^+ form) and PD-10 columns were obtained from GE Healthcare Life-Sciences (Little Chalfont, UK). Reactive Red 120-Agarose and Dowex 50W-X4 (200 mesh) resins came from Sigma-Aldrich, and Vivaspin-500 and 20 centrifugal concentrators were from Sartorius Stedim (Goettingen, Germany). All other enzymes and DNA modifying enzymes as well as the TurboFect transfection reagent were obtained from Thermo-Fermentas (Waltham, USA), A&A Biotechnology (Gdynia, Poland) or BioShop (Burlington, Canada).

### Assay of the Actin-specific Histidine *N*-methyltransferases Activity

#### Protein Substrates

The enzyme activity was determined by measuring the incorporation of [^3^H]methyl group from S-[methyl-^3^H]adenosyl-L-methionine ([^3^H]SAM) into homogenous recombinant human (mammalian) β-actin or its mutated form in which histidine 73 was replaced by alanine residue (H73A). The standard incubation mixture (0.11 ml) contained 25 mM Tris-HCl, pH 7.2, 10 mM KCl, 1 mM DTT, 2 μM protein substrate and 1 μM [^1^H+^3^H] SAM (≈ 400 × 10^3^ cpm). When appropriate, the incubation mixture was supplemented with recombinant *S*-adenosyl-L-homocysteine (SAH) nucleosidase and adenine deaminase as indicated in legends to figures and tables. The reaction was started by the addition of enzyme preparation and carried out at 37°C for 15 min unless otherwise described. Protein methylation was linear for at least 15 min under all conditions studied. By analogy to assays of nonribosomal peptide synthetase activity [Richardt *et al.*, 2003], the incubation was stopped by the addition of 0.1 ml of the reaction mixture to 0.025 ml of BSA (1 mg/ml) and 0.8 ml of ice-cold 10 % (w/v) trichloroacetic acid (TCA). After 30 min on ice, the precipitate was pelleted and washed twice with ice-cold 10 % TCA. The pellet was finally dissolved in pure formic acid.

#### Peptide Substrates

The enzyme-dependent methylation of synthetic peptides (Peptide H: YPIEHGIVT and Peptide A:YPIEAGIVT), analogues of methylation sites in protein substrates, was determined by measuring the incorporation of [^3^H]methyl group from [^3^H]SAM into a corresponding peptide. The composition of a standard incubation mixture was similar to that described above, excepting that protein substrates were replaced by peptides at 2 mM concentration. Blanks containing no peptides were included in all assays. The reaction was started by the addition of enzyme preparation and carried out at 37°C for time periods indicated in the figure caption. The incubation was stopped by the addition of 0.1 ml of the reaction medium to 0.2 ml of ice-cold 10 % (w/v) HClO_4_. The samples were diluted with 0.12 ml of H_2_O and centrifuged at 13,000 × g for 10 min. After neutralization of the supernatant with 3 M K_2_CO_3_, the salts were removed by centrifugation (13,000 × g for 10 min) and the clear supernatant was diluted 5 times with 20 mM Hepes, pH 7.5 and 2 ml were applied to Dowex 50W-X4 columns (1 ml, Na^+^ form), equilibrated with 20 mM Hepes, pH 7.5. The columns were washed with 5 × 2 ml of 20 mM Hepes pH 7.5, and resulting fractions contained methylated peptides. To elute a non-consumed [^1^H + ^3^H]SAM, the columns were washed with 4 × 2 ml of 1 M NH_4_OH.

In all cases, samples to be counted were mixed with 10 ml of scintillation fluid (Ultima Gold) and the incorporated radioactivity was analyzed with a Beckman LS6000 IC liquid scintillation counter.

### Protein Purification

#### Purification of Rat Actin-specific Histidine N-methyltransferase

A myofibrillar extract of rat skeletal muscle was prepared as described by Vijayasarathy and Rao [1987]. Briefly, rat leg muscles (200 g) from 18 male WAG rats, aged 3 months, were homogenized with 3 volumes (w/v) of buffer consisted of 10 mM Tris-HCl, pH 7.4, 1 mM DTT, 250 mM sucrose, 5 μg/ml leupeptin and 5 μg/ml antipain, with a Waring Blender 7011HS. The homogenate was centrifuged for 15 min at 10,000 × g at 4°C. The pellet was resuspended in 2400 ml of borate buffer (39 mM potassium borate, pH 7.2, 275 mM KCl, 1 mM DTT, 3 μg/ml leupeptin and 3 μg/ml antipain), extracted with shaking at 160 rpm for 1h at 5°C and centrifuged for 15 min at 10,000 × g at 4°C. The resulting supernatant solution (2400 ml) was then fractionated between 0% and 20% concentration (w/v) of PEG 4000. The 0-20% precipitate was dissolved in 1000 ml of buffer A (20 mM Tris-HCl, pH 8.0, 1 mM DTT, 3 μg/ml leupeptin and 3 μg/ml antipain), centrifuged for 20 min at 25,000 × g at 4°C and filtered through 6 layers of gauze to remove fat particles.

The clarified supernatant was applied to a DEAE-Sepharose column (350 ml bed volume) equilibrated with buffer A. The column was washed with 500 ml of buffer A, developed with a NaCl gradient (0-1 M in 962 ml) in buffer A and fractions (6.5 ml) were collected. The enzymatically active fractions of the DEAE-Sepharose column (84 ml) were diluted to 350 ml with buffer A and applied to a Q-Sepharose column (100 ml bed volume) equilibrated with the same buffer. The column was washed with 360 ml of buffer A and the retained protein was eluted with a NaCl gradient (0-0.95 M in 500 ml in buffer A) and fractions (5 ml) were collected. The active fractions of the Q-Sepharose column (64 ml) were pooled, supplemented with solid NaCl to reach the final salt concentration in the sample equal to 0.6 M, and applied to a Phenyl-Sepharose 6 column (40 ml bed volume) equilibrated with buffer B (20 mM Tris-HCl, pH 7.2, 10 mM KCl, 1 mM DTT, 2 μg/ml leupeptin and 2 μg/ml antipain) containing 0.6 M NaCl. The hydrophobic interaction column was washed with 100 ml of buffer B with NaCl and the retained protein was eluted with a NaCl gradient (600-0 mM in 360 ml in buffer B). Fractions (5 ml) were collected. The enzymatically active fractions of the Phenyl-Sepharose column (93 ml) were pooled and loaded on a HiScreen Blue-Sepharose column (4.7 ml bed volume) equilibrated with buffer B. The column was washed with 20 ml of buffer B, developed with a NaCl gradient (0-1.5 M in 70 ml) in buffer B and fractions (3 ml) were collected. The active fractions of the HiScreen Blue-Sepharose column (27 ml) were pooled, supplemented with 100 mM KCl, 2 mM ATP, 0.2 mM CaCl_2_ and concentrated to 2.5 ml in Vivaspin-20 ultrafiltration devices. Two milliliters of the ultrafiltrate were loaded on a Superdex 200 16/60 column (120 ml bed volume) equilibrated with buffer C (20 mM Tris-HCl, pH 7.2, 100 mM KCl, 1 mM DTT, 0.2 mM ATP, 0.2 mM CaCl_2_, 2 μg/ml leupeptin and 2 μg/ml antipain). The gel filtration column was then developed with 140 ml of buffer C and 3 ml fractions were collected. To obtain more purified enzyme preparation for a tandem mass spectrometry, the most active fraction (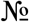 14, 2.5 ml) of Superdex 200 purification step was loaded on a Reactive Red 120-Agarose column (1.5 ml bed volume) equilibrated with buffer C. The column was first washed with 6 ml of buffer C and the retained proteins were eluted (4 fractions of 3 ml) with buffer C containing an increasing concentration of NaCl (0-1.4 M). All purification steps were performed at 4°C and the enzymatic preparation was stored at -70°C between steps.

#### Overexpression and Purification of the Recombinant β-actin Inclusion-body Protein

Plasmid pCOLD I encoding human β-actin (ACTB, GenBank: NM_001101.4) was a kind gift of Dr. Minoru Tamura (Ehime University, Japan) and was prepared as described in [Tamura *et al.*, 2011].

For β-actin production, Escherichia coli BL21(DE3) (Agilent, USA) cells were transformed with the DNA construct and a single colony was selected to start an over-nigh pre-culture. 500 mL of LB broth (with 100 μg/mL ampicilin) was inoculated with 50 ml of the pre-culture and incubated at 37 °C and 200 rpm until an OD600 of 0.5 was reached. The culture was placed on ice for 20 min. (cold-shock) and IPTG was added to a final concentration of 0.2 mM to induce protein expression. Cells were incubated for 20 h at 15 °C, 200 rpm, and harvested by centrifugation (6000 × g for 10 min). The cell paste was resuspended in 27.5 ml lysis buffer consisted of 20 mM Hepes, pH 7.5, 1 mM DTT, 1 mM ADP, 0.5 mM PMSF, 2 μg/ml leupeptin and 2 μg/ml antipain, 0.2 mg/ml hen egg white lysozyme (BioShop), 1000 U Viscolase (A&A Biotechnology). The cells were lysed by freezing in liquid nitrogen and after thawing and vortexing, the extracts were centrifuged at 4° C (20,000 × g for 30 min).

The pellet, containing inclusion bodies, was completely resuspended in buffer A (20 mM Hepes, pH 7.5, 2M urea, 0.5 M NaCl, 5 mM DTT, 2 mM EDTA) with the use of Potter-Elvehjem homogenizer and centrifuged at 4° C (20,000 × g for 10 min). The resulting pellet was then subjected to a further two rounds of sequential wash in buffer B (20 mM Hepes, pH 7.5, 0.5 M NaCl, 5 mM DTT, 2 mM EDTA) and buffer C (20 mM Hepes, pH 7.5, 0.5 M NaCl). The washed inclusion bodies were finally solubilized in loading buffer (20 mM Tris-HCl, pH 7.5, 6 M guanidine HCl, 0.5 M NaCl, 10 mM imidazole) and applied on a HisTrap FF column (5 ml) equilibrated with the same buffer.

The column was washed with 20 ml of loading buffer and the bound protein was refolded by the use of a linear gradient of 6-0 M guanidine HCl in loading buffer (40 ml for 20 min). Next, the column was washed with 15 ml of loading buffer without guanidine HCl and the retained proteins were eluted with a stepwise gradient of imidazole (25 ml of 40 mM, 25 ml of 60 mM and 20 ml of 500 mM). The recombinant proteins were eluted with 500 mM imidazole in homogeneous form as confirmed by SDS-PAGE (Fig. 1). The β-actin preparation was immediately desalted on PD-10 columns equilibrated with 20 mM Tris-HCl, pH 7.5, 1 mM DTT, 6% sucrose, 2 μg/ml leupeptin and 2 μg/ml antipain. The purified β-actin was stored at -70°C.

**Figure 1.**
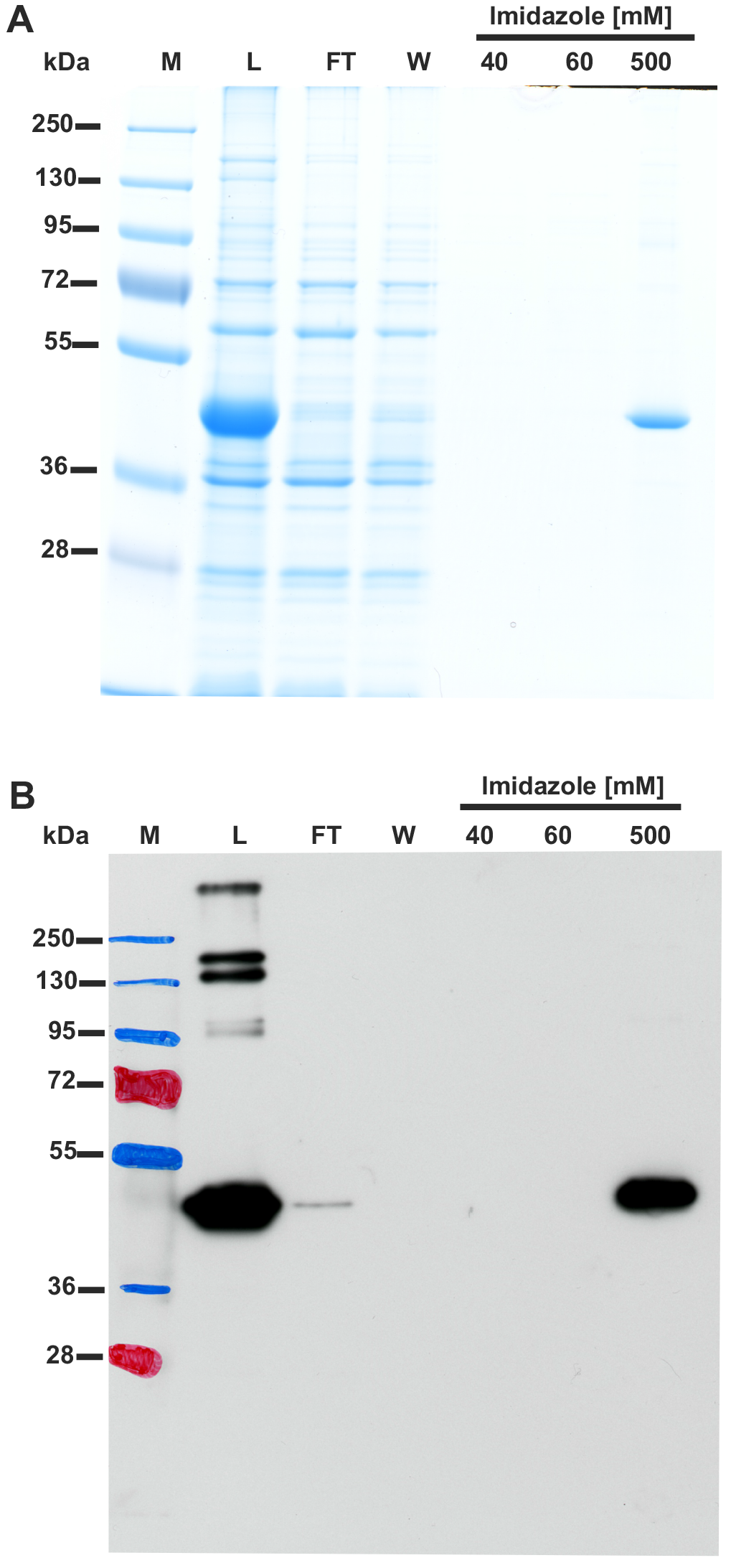
SDS-PAGE (A) and Western-blot (B) analysis of fractions obtained during purification of recombinant human β-actin overexpressed in *E. coli*. Human β-actin was purified to homogeneity by the affinity chromatography on nickel-Sepharose (HisTrap HP) as described under “Experimental Procedures”. For the SDS-PAGE analysis, 20 μl of sample from each fraction was loaded onto a 10 % gel, electrophoresed and the resulting gel was then stained with colloidal Coomassie blue. For the Western-blot analysis, 1 μl of each fraction was loaded onto a 10 % gel, electrophoresed and blotted to nitrocellulose membrane which was then sequentially probed with a mouse primary antibody against His_6_ tag and a horseradish peroxidase-conjugated goat anti-mouse antibody. Secondary antibody was detected through autoradiography using chemiluminescence. M, prestained protein marker; L, urea-washed inclusion bodies of *E. coli* applied on the column; FT, flow through; W, wash; Fractions 40 to 500 were eluted with the indicated concentrations of imidazole.

#### Overexpression and Purification of the Soluble Recombinant β-actin

Plasmid pCOLD I encoding human β-actin was used as a template for PCR amplification of β-actin cDNA. The resulting ORF was then cloned into pESC-URA yeast expression vector. *Saccharomyces cerevisiae* BY4742 strain (EUROSCARF, Germany) was transformed with above-mentioned plasmid using high-efficiency lithium acetate transformation method [Gietz and Woods, 2002]. Resulting yeast strain was designated as BY4742/β-actin.

Method for the purification of recombinant β-actin expressed in *S. cerevisiae* was as reported by Karlsson [1988]. BY4742/β-actin was cultivated in 500 ml of SG medium (0,67% yeast nitrogen base without amino acids, 2% galactose, supplemented with histidine, leucine and lysine) for 24 h, cells were collected, washed with water, and suspended in 30 ml of buffer C (5 mM Tris-HCl pH 7.6, 0.2 mM CaCl_2_, 0.2 mM DTT, 0.3 mM PMSF). After addition of 100 U of DNase I, cells were homogenized in Braun MSK cell homogenizer using glass beads (B. Braun Melsungen). Crude lysate was centrifuged (5000 × g, 5 min), diluted twice with the buffer, and applied to the Ni-NTA-agarose column (Sigma-Aldrich). Next, the column was washed with 30 ml of buffer A supplemented with 8 mM imidazole, and retained proteins were eluted with buffer A containing 500 mM imidazole (≈15 ml, 0.7 ml fractions). Protein concentration in each fraction was estimated by SDS-PAGE and the most concentrated samples were taken for further procedure.

For preparation of β-actin complexes with either cofilin or profilin (see below), equimolar amount of the recombinant homogenous actin-binding protein was mixed with the yeast-producing β-actin and dialyzed twice against 1000 ml of dialysis buffer (20 mM Tris-HCl pH 7,6, 1 mM DTT, 6% sucrose, 0.2 mM CaCl_2_), for 2 h in 250 ml and then overnight in 500 ml. When obtaining β-actin complexed with either ADP or ATP, samples eluted from Ni-NTA-agarose were dialyzed under the same conditions against dialysis buffer containing 50 μM of an appropriate nucleotide. In all cases, dialyzed protein preparation was concentrated using Amicon 10K device (Merck,).

#### Overexpression and Purification of the Recombinant SETD3 Proteins

Rat total RNA was prepared from 200 mg of leg muscles with the use of TriPure reagent according to the manufacturer’s instructions, while human skeletal muscle total RNA was purchased from Clontech (USA). cDNA was synthesized using Moloney murine leukemia virus-reverse transcriptase (Thermo-Fermentas), with oligo(dT)_18_ primer and 2.5 μg total RNA according to the manufacturer’s instructions.

The open reading frames encoding rat (NCBI Reference Sequence: XM_002726774.2) and human SETD3 protein (NM_032233.2) were PCR-amplified using Pfu DNA polymerase in the presence of 1 M betaine. SETD3s were amplified using 5’ primers containing the initiator codon preceded by the Kozak consensus sequence [Kozak, 1987] and an KpnI site and 3’ primers in which the original stop codon was replaced by an amino-acid coding codon flanked by a NotI site (for primer sequences, see Table 1). The amplified DNA products of the expected size were digested with the appropriate restriction enzymes and cloned in the pEF6/Myc-His A expression vector (Invitrogen, USA), which allows the production of proteins with an C-terminal His_6_ tag, and verified by DNA sequencing (Macrogen, Netherlands). For transfections, COS-7 cells (Cell Lines Service, Germany) or HEK-293T (a kind gift of Dr. Maria Veiga-da-Cunha, de Duve Institute, Belgium) were plated in 100-mm Petri dishes at a cell density of 1.7 × 10^6^ or 2.1 × 10^6^ cells per plate, respectively, in Dulbecco’s minimal essential medium supplemented with 100 units/ml penicillin, 100 μg/ml streptomycin, and 10% (v/v) fetal bovine serum, and grown in a humidified incubator under 95% air and 5% CO_2_ atmosphere at 37°C. After 24 h, each plate was transfected with 7-8 μg of either unmodified pEF6/Myc-His A vector or the same vector encoding SETD3 protein using the TurboFect transfection reagent according to the protocol provided by the manufacturer. After 48h the culture medium was removed, the cells were washed with 5 ml phosphate buffered saline and harvested in 1 ml of 20 mM Tris-HCl, pH 7.2, containing 10 mM KCl, 1 mM DTT, 5 μg/ml leupeptin and 5 μg/ml antipain. The cells were lysed by freezing in liquid nitrogen and after thawing and vortexing, the extracts were centrifuged at 4° C (20,000 × g for 30 min) to remove insoluble material.

**Table 1.**
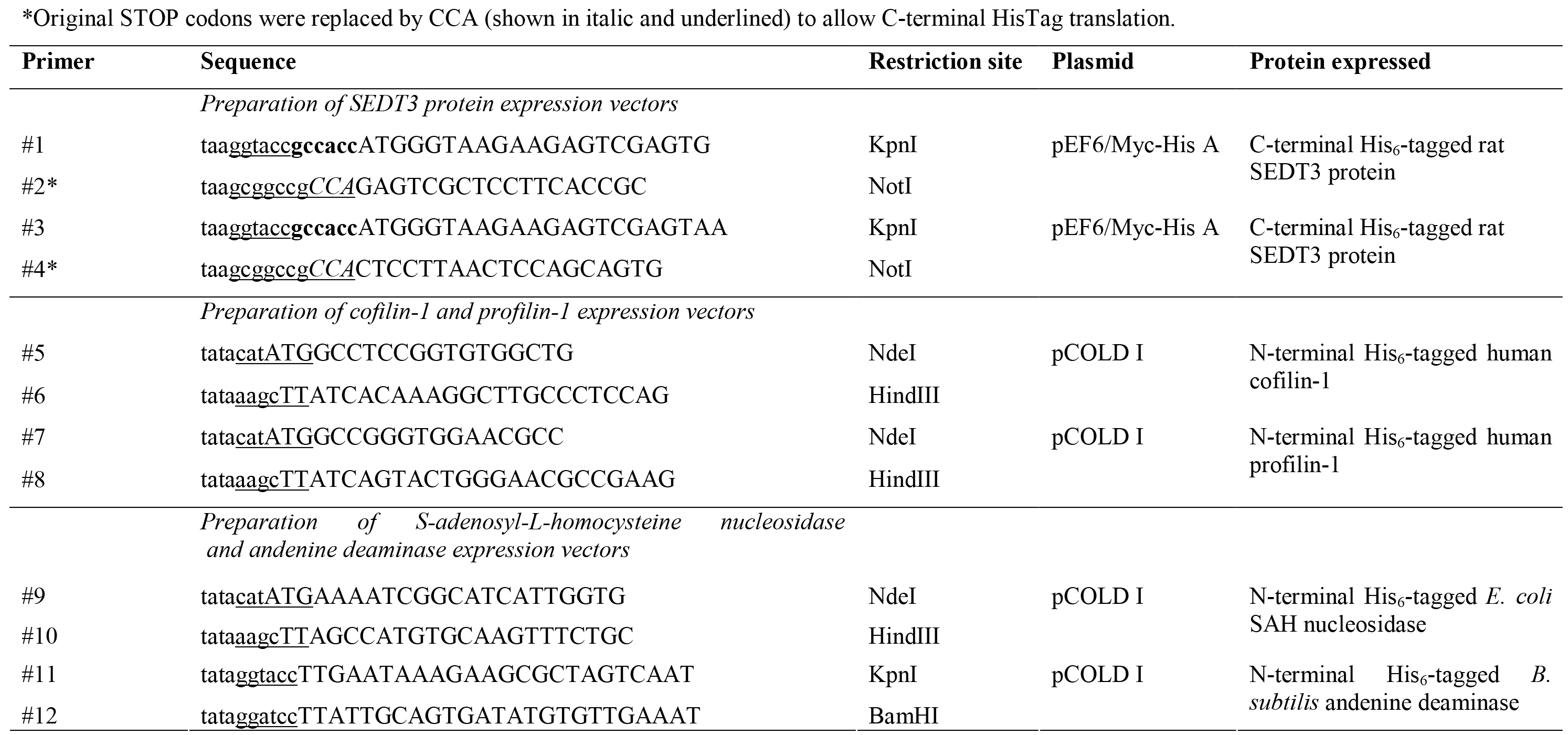
Sequences of primers used in PCR experiments. The nucleotides corresponding to the coding sequences are in capital letters, the Kozak consensus sequence is shown in bold, and the restriction sites added are underlined.

For the purification of recombinant SETD3 proteins, the supernatant of COS-7 lysate (13-14 ml) was diluted 3-fold with buffer A (50 mM Tris-HCl, pH 7.4, 400 mM NaCl, 10 mM KCl, 30 mM imidazole, 3 μg/ml leupeptin and 3 μg/ml antipain) and applied on a HisTrap HP column (1 ml) equilibrated with the same buffer. The column was washed with 6-10 ml buffer A and the retained protein was eluted with a stepwise gradient of imidazole (6 ml of 60 mM, 6 ml of 150 mM and 6 ml of 300 mM) in buffer A. The recombinant proteins were eluted with 150 mM imidazole in homogeneous form as confirmed by SDS-PAGE (Fig. 2). The enzyme preparations were desalted on PD-10 columns equilibrated with 20 mM Tris-HCl, pH 7.2, 50 mM KCl, 1 mM DTT, 6% sucrose, 2 μg/ml leupeptin and 2 μg/ml antipain. The yield of recombinant proteins was 0.5-0.6 mg of homogenous protein per about 25-40 mg of soluble COS-7 cell protein. The purified enzymes were aliquoted and stored at -70°C. Repeated freeze-thawing was avoided, since it impairs SETD3 activity.

**Figure 2.**
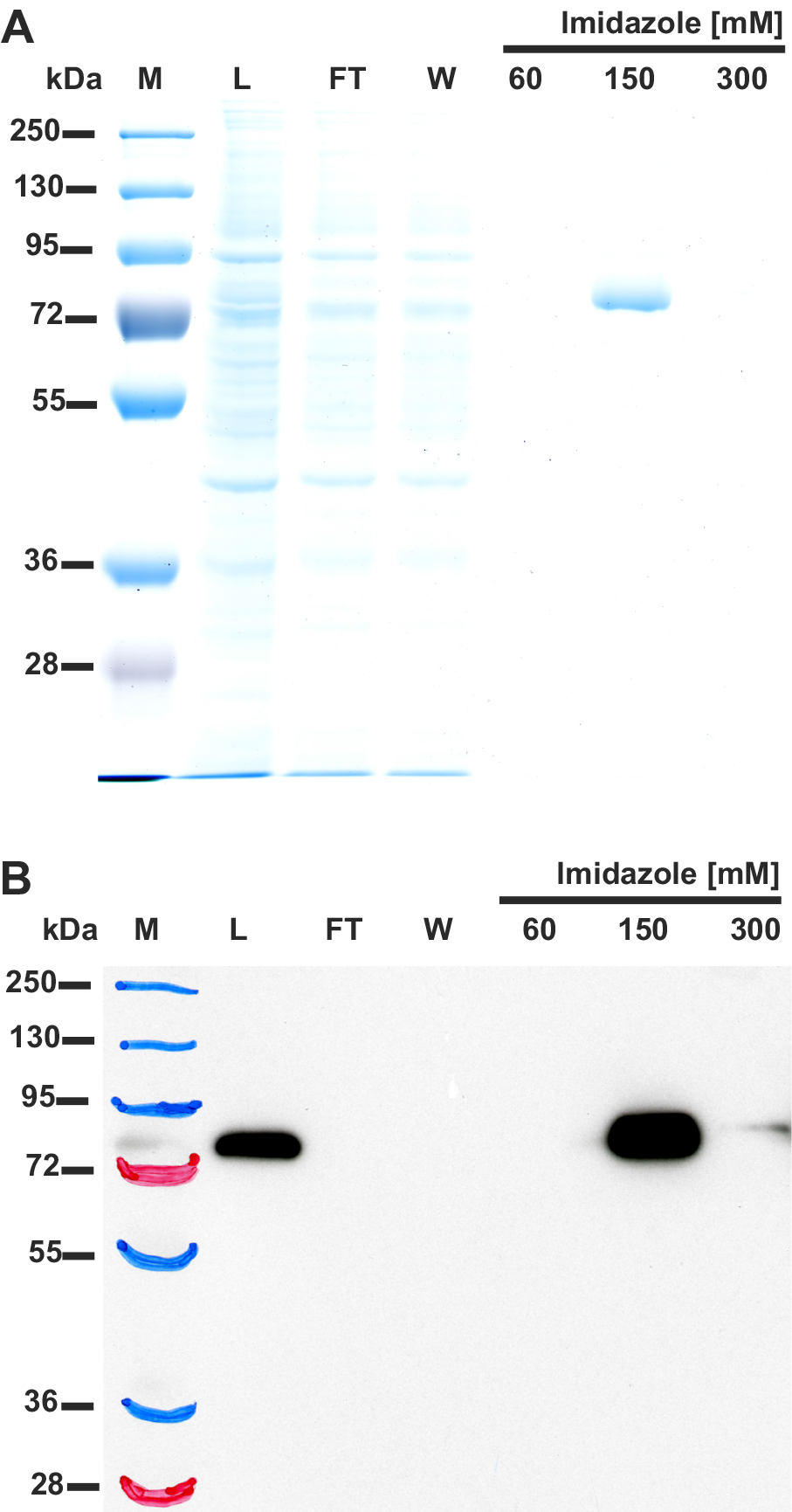
SDS-PAGE (A) and Western-blot (B) analysis of fractions obtained during purification of the recombinant rat SETD3 protein. Rat SETD3 protein was purified to homogeneity by affinity chromatography on the nickel-Sepharose (HisTrap HP) as described under “Experimental Procedures”. For the SDS-PAGE analysis, 15 μl of sample from each fraction was loaded onto a 10% gel, electrophoresed and the resulting gel was then stained with silver [Shevchenko *et al.*, 1996]. For the Western-blot analysis, 7.5 μl of each fraction was loaded onto a 10% gel, electrophoresed and blotted to nitrocellulose membrane which was then sequentially probed with a mouse primary antibody against His6 tag and a horseradish peroxidase-conjugated goat anti-mouse antibody. Secondary antibody was detected using enhanced chemiluminescence. M, prestained protein marker; L, cell-free lysate of COS-7 cells applied on the column; FT, flow through; W, wash; Fractions 60 to 300 were eluted with the indicated concentrations of imidazole.

#### Overexpression and Purification of the Recombinant Cofilin and Profilin Proteins

Human skeletal muscle total RNA was from Clontech (USA) and cDNA was synthesized as described above. The open reading frames encoding human cofiline-1 and profiline-1 (NCBI Reference Sequence: NM_005507.2 and NM_005022.3, respectively) were PCR-amplified using Pfu DNA polymerase in the presence of 1 M betaine. ORFs of both proteins were amplified using 5’ primers containing the initiator codon preceded by an NdeI site and 3’ primers flanked by a HindIII site (for primer sequences, see Table 1). The amplified DNA products of the expected size were digested with the appropriate restriction enzymes and cloned into the pCOLD I expression vector (Takara Bio, Kusatsu, Japan), which allows the production of proteins with an N-terminal His_6_ tag, and verified by DNA sequencing (Macrogen, Netherlands).

For protein production, *E. coli* BL21(DE3) cells were transformed with the appropriate DNA construct and a single colony was selected to start an over-nigh pre-culture. 200 mL of LB broth (with 100 μg/mL ampicilin) was inoculated with 20 ml of the pre-culture and incubated at 37 °C and 200 rpm until an OD600 of 0.6 was reached. The culture was placed on ice for 20 min. (cold-shock) and IPTG was added to a final concentration of 0.25 mM to induce protein expression. Cells were incubated for 16 h at 13°C, 200 rpm, and harvested by centrifugation (6000 × g for 10 min). The cell paste was resuspended in 10 ml lysis buffer consisted of 25 mM Hepes, pH 7.5, 300 mM NaCl, 50 mM KCl, 1 mM DTT, 2 mM MgCl_2_, 1 mM PMSF, 5 μg/ml leupeptin and 5 μg/ml antipain, 0.25 mg/ml hen egg white lysozyme (BioShop, Canada), 50 U DNase I (Roche). The cells were lysed by freezing in liquid nitrogen and after thawing and vortexing, the extracts were centrifuged at 4° C (20,000 × g for 30 min).

For the purification of recombinant actin-binding proteins, the supernatant of *E. coli* lysate (10 ml) was diluted 4-fold with buffer A (50 mM Hepes, pH 7.5, 300 mM NaCl, 10 mM KCl, 30 mM imidazole, 1 mM DTT) and applied on a HisTrap FF column (5 ml) equilibrated with the same buffer. The column was sequentially washed with 23 ml buffer A and 23 ml of the same buffer supplemented with urea at 1.8 M concentration. Next, the retained protein was eluted with a stepwise gradient of imidazole (23 ml of 60 mM, 23 ml of 150 mM and 22 ml of 300 mM) in buffer A. The recombinant proteins were eluted with 300 mM imidazole in homogeneous form as confirmed by SDS-PAGE (not shown). The enzyme preparations were dialyzed three times against 400 ml of dialysis buffer consisted of 20 mM Tris-HCl, pH 7.5, 200 mM NaCl, 1 mM DTT, 6% sucrose. The yield of recombinant proteins was 7.1 and 17.7 mg of homogenous cofilin-1 and profilin-1, respectively, per 200 ml of *E. coli* culture. The purified proteins were aliquoted and stored at -70°C.

#### Overexpression and Purification of the Recombinant SAH Nucleosidase and Adenine Deaminase

*E. coli* DNA was extracted by heating 50 μl of over-night culture of BL21(DE3) cells at 95°C for 15 min, whereas *Bacillus subtilis* (ATCC 6633, Sigma-Aldrich) genomic DNA was purified from 100 mg of bacterial cells with the use of TriPure reagent according to the manufacturer’s instructions.

The open reading frames encoding *E. coli* SAH nucleosidase and *B. subtilis* adenine deaminase (NCBI Reference Sequence: NC_000913.3 and NC_000964.3, respectively) were PCR-amplified using Pfu DNA polymerase solely or mixture of Taq: Pfu polymerases (1:0.2), respectively, in the presence of 1 M betaine. SAH nucelosidase ORF was amplified using 5’ primer containing the initiator codon preceded by an NdeI site and 3’ primer with a HindIII site, while for the amplification of adenine deaminase DNA, 5’ primer with the initiator codon preceded by an KpnI site and 3’ primer with a BamHI site were used (for primer sequences, see Table 1). The amplified DNA products of expected size were digested with the appropriate restriction enzymes, cloned into the pCOLD I expression vector and verified by DNA sequencing (Macrogen, Amsterdam, Netherlands).

For protein production, *E. coli* BL21(DE3) cells were transformed with the appropriate DNA construct and a single colony was selected to start an over-nigh pre-culture. 100 mL of LB broth (with 100 μg/mL ampicilin) was inoculated with 10 ml of the pre-culture and incubated at 37 °C and 200 rpm until an OD600 of 0.6 was reached. The culture was placed on ice for 20 min (cold-shock) and IPTG was added to a final concentration of 0.25 mM to induce protein expression. Cells were incubated for 16 h at 13°C, 200 rpm, and harvested by centrifugation (6000 × g for 10 min). The cell pellet was resuspended in 10 ml lysis buffer consisted of 25 mM Hepes, pH 7.5, 300 mM NaCl, 50 mM KCl, 1 mM DTT, 2 mM MgCl_2_, 1 mM PMSF, 5 μg/ml leupeptin and 5 μg/ml antipain, 0.25 mg/ml hen egg white lysozyme (BioShop, Canada), 25 U DNase I (Roche). The cells were lysed by freezing in liquid nitrogen and after thawing and vortexing, the extracts were centrifuged at 4°C (20,000 × g for 30 min).

For protein purification, the supernatant of *E. coli* lysate (10 ml) was diluted 3-fold with buffer A (50 mM Tris-HCl, pH 7.2, 400 mM NaCl, 10 mM KCl, 30 mM imidazole, 1 mM DTT, 3 μg/ml leupeptin and 3 μg/ml antipain) and applied on a HisTrap FF column (5 ml) equilibrated with the same buffer. The column was washed with 20-30 ml buffer A and the retained protein was eluted with a stepwise gradient of imidazole (25 ml of 60 mM, 20 ml of 150 mM and 20 ml of 300 mM) in buffer A. The recombinant proteins were eluted with 150-300 mM imidazole in homogeneous form as confirmed by SDS-PAGE (not shown). The enzyme preparations were desalted on PD-10 columns equilibrated with 20 mM Tris-HCl, pH 7.2, 50 mM KCl, 1 mM DTT, 6% sucrose, 2 μg/ml leupeptin and 2 μg/ml antipain. The yield of recombinant proteins was 1.2 and 3.1 mg of homogenous adenine deaminase and SAH nucleosidase, respectively, per 200 ml of *E. coli* culture. The purified enzymes were aliquoted and stored at -70°C.

### Identification of the Rat Actin-specific Histidine *N*-methyltransferase by Tandem Mass Spectrometry

Since a preliminary SDS-PAGE analysis of the peak activity fractions from the Reactive Red 120-agarose purification step revealed the presence of very faint protein bands with a low protein content, the fractions were 15-fold concentrated in a Vivaspin-500 ultrafiltration device and reanalyzed by SDS-PAGE. All bands present in the concentrated fractions were cut from a 10% gel and digested with trypsin. In-gel digestions of the peptides were performed as described in [Shevchenko *et al.*, 2006]. Peptides were analyzed by nanoUPLC-tandem mass spectrometry employing Acquity nanoUPLC coupled with Synapt G2 HDMS Q-TOF mass spectrometer (Waters, Milford, USA) fitted with a nanospray source and working in MS^E mode under default parameters. Briefly, products of in-gel protein digestion were loaded onto a Waters Symmetry C18 trapping column (20 mm × 180 μm) coupled to the Waters BEH130 C18 UPLC column (250 mm × 75 μm). The peptides were eluted from these columns in a 1-85% gradient of acetonitrile in water (both containing 0.1% formic acid) at a flow rate of 0.3 μl/min. The peptides were directly eluted into the mass spectrometer. Data were acquired and analyzed using MassLynx 4.1 software (Waters, USA) and ProteinLynx Global Server 2.4 software (PLGS, Waters, USA) with a False Discovery Rate ≤ 4 %, respectively. To identify actin-specific histidine *N*-methyltransferase, complete rat (*Rattus norvegicus*) reference proteome was downloaded from the NCBI Protein database, randomized and used as a databank of the MS/MS software.

### Generation of the HAP1 Cell Lines Deficient in SEDT3 by CRISPR/Cas9 Method

The human-derived HAP1 cell line (Horizon Discovery, Waterbeach, UK) expresses *Setd3* gene at the level of 32.04 transcripts per kilobase million (https://www.horizondiscovery.com/human-setd3-knockout-cell-line-hzghc84193). The parental cells were therefore used to create HAP1 cell lines deficient in SETD3 by CRISPR/Cas9 technology (DNA double nicking strategy).

Two CRISPR/Cas9 constructs were generated to target exon 4 of human *Setd3* gene by ligating two different sets of annealed primer pairs (CRISPR-hSETD3-s1 CACCGgttctgtcgagggttttgaaa, CRISPR-hSETD3-as1 AAACtttcaaaaccctcgacagaacC and CRISPR-hSETD3-s2 CACCGgaggcccatttcattagatc, CRISPR-hSETD3-as2 AAACgatctaatgaaatgggcctcC). The annealed primer pairs were ligated into the vector pSpCas9n(BB)-2A-Puro (PX462) V2.0 (a gift from F. Zhang, Massachusetts Institute of Technology; Addgene plasmid no. 62987) [Ran *et al.*, 2013] digested by BbsI. Constructs were validated by sequencing (Beckman Coulter Genomics). HAP1 cells were cultured in IMDM (Iscove’s Modified Dulbecco’s Medium) containing 10% FBS, 2 mM L-glutamine and penicillin/streptomycin (Thermo-Life Technologies). Cells were transfected with the CRISPR constructs exactly as described in [Collard *et al.*, 2016]. Genomic DNA from puromycin resistant clones was used to PCR-amplify the regions encompassing the targeted site and the PCR products were sequenced to evaluate the gene modification present in each clone. The three SETD3-KO clones that were ultimately selected for further studies presented the following modifications: clone KO-A1, a 16-bp deletion after Phe71, that introduces a frame shift and a premature stop codon at position 75; clone KO-A3, a 4-bp deletion and 6-bp insertion introducing a frame shift after Met83 and a premature stop codon at position 109; clone KO-A5, a 18-bp deletion and 107-bp insertion after Trp85, with a premature stop codon at the position 292.

The absence of SETD3 protein in both SETD3-KO cell lines was confirmed by western blotting performed as described previously [Drozak *et al.*, 2013] with the use of rabbit polyclonal primary antibody against human SETD3 (ab174662, Abcam, UK) and a horseradish peroxidase-conjugated goat anti-rabbit secondary IgG antibody (Agrisera, Sweden).

### Product Analysis

#### Detection of In Vivo Methylated β-actin by Tandem Mass Spectrometry

HAP1 wild-type (WT-A1 and WT-A2) and SETD3-KO cells (KO-A1 and KO-A3) were cultured in IMDM containing 10% FBS, 2 mM L-glutamine and penicillin/streptomycin (Life Technologies), each in four 10-cm diameter plates. At confluence, the medium was rapidly removed, the plates were washed with 5 ml cold PBS and each off them scraped in 0.8 ml of buffer A (25 mM Hepes, pH 7.4, 20 mM NaCl, 2.5 μg/ml leupeptin and 2.5 μg/ml antipain) and combined to collect the cells. The four soluble cell extracts were prepared by 2 cycles of freezing the cell suspension in liquid nitrogen/thawing and vortex-mixing and centrifuged for 15 min at 15,000 × g and 4 °C to recover soluble proteins.

To partially purify the actin, the cell lysates were diluted 5-fold in the buffer A (20 ml final volume) and loaded onto a 1 ml-HiTrap Q HP (GE Healthcare) equilibrated with the same buffer. The column was washed with 6 ml of buffer A and proteins were eluted with a linear NaCl gradient (20-750 mM in 20 ml buffer A). The elution fractions (1 ml) were analysed by SDS-PAGE followed by Coomassie blue staining and by western blotting using a rabbit anti-actin (A2066, Sigma-Aldrich). The actin band of the richest fraction was cut out of the gel and digested with trypsin.

The resulting peptides were analysed by LC-MS/MS with a LTQ XL (ThermoFisher Scientific) equipped with a microflow ESI source interfaced to a Dionex Ultimate Plus Dual gradient pump, a Switchos column switching device, and Famos Autosampler (Dionex, Amsterdam, Netherlands) as described (Houddane *et al.*, 2017). Raw data files were then analysed by bioinformatics using Proteome Discoverer (Thermo Fisher Scientific). Peak lists were generated using the application spectrum selector in the Proteome Discoverer 1.4 package. From raw files, MS/MS spectra were exported as individual files in .dta format with the following settings: peptide mass range: 400-3500 Da; minimal total ion intensity: 500; minimal number of fragment ions: 12. The resulting peak lists were searched using SequestHT against a human protein database obtained from Uniprot and containing only actin sequences. The following parameters were used: trypsin was selected with semi-proteolytic cleavage only after lysine and arginine; the number of internal cleavage sites was set to 2; the mass tolerance for precursor and fragment ions was 1.0 Da; and the considered dynamic modifications were +15.99 Da for oxidized methionine, +14.00 Da for methylation on histidine and +42.00 Da for acetylation of N-terminus or lysines. Peptide spectral matches (PSM) were filtered using charge-state versus cross-correlation scores (Xcorr) and methylation sites were validated manually.

#### Detection of In Vitro Methylated β-actin by Tandem Mass Spectrometry

To obtain sufficient amount of the methylated β-actin formed in the reaction catalyzed by recombinant rat or human SETD3 protein for mass spectrometry analysis, the reaction mixture was scaled up. Briefly, 15 μg recombinant human β-actin was incubated for 90 min at 37°C in 70 μl of a reaction mixture containing 25 mM Tris-HCl, pH 7.2, 10 mM KCl, 1 mM DTT, 20 μM MnCl_2_, 6% sucrose, 20-30 μM ([^2^H]SAM), 4.5-9.0 μg SAH nucleosidase and 1.75-3.5 μg adenine deaminase in the absence or presence of 1.1-3.5 μg SETD3 protein. The incubation was stopped by the addition of 30 μl of the reaction mixture to 10 μl 4× Laemmli Buffer and heating the sample for 5 min at 95°C, while the reaming 40 μl of the mixture was flesh-frozen in LN2 and stored at -70°C. The yield of β-actin methylation was about 30% as determined for the parallel labeling reaction in the presence of [^1^H +^3^H]SAM.

For MS/MS analysis of deuterated protein, 35 μl of denatured sample (≈18 μg protein) were separated in 10% SDS-PAGE and silver stained [Shevchenko *et al.* 1996]. The bands corresponding to β-actin were excised from the gels, destained and digested with trypsin according to [Shevchenko *et al.* 2006]. Additionally, 18 μl of the flesh-frozen sample (12 μg protein) were trypsin-digested “in solution” for 16 h at 30°C in 30 μl of the reaction mixture containing 100 mM NH_4_HCO_3_ and 1.5 μg trypsin (MS grade, Sigma-Aldrich). The digestion reaction was terminated by the addition of trifluoroacetic acid to a final concentration of 1%. Resulting peptides were analyzed by nanoUPLC-tandem mass spectrometry as described above in the section “*Identification of Rat Actin-specific Histidine N-methyltransferase by Tandem Mass Spectrometry*”. To identify actin peptides, the full amino acid sequence of recombinant human β-actin in fusion with His_6_-tag at N-terminus was used as a databank of the MS/MS software. Detection of the trideuterium-methylated peptides was performed in a fully automatic mode by PLGS 2.4 software (Waters, USA), following its update by adding trideuterium-methylation (+17,03448 Da) of Cys, Asp, Asn, His, Lys, Arg, Glu, Gln residues as a possible modification of peptides.

### Phenotypic Analysis

#### Lactate Production

HAP1 cells (150 × 10^3^) were seeded in 12-well plates and incubated in IMDM with 10% FBS for 40 h with a change of medium every 20 hours. Following 40 h incubation, cell confluency was about 90% and the fresh medium was added to start the final incubation lasting for next 20 h. To finish the incubation, 1 ml of culture medium was transferred to 100 μl of 35% perchloric acid (PCA), centrifuged at 10,000 ×g for 10 min, and resulting supernatant was neutralized with 3 M K_2_CO_3_. Lactate concentration was determined spectrophotometrically using lactate dehydrogenase (Roche, Poland) according to Noll [1983]. Adherent cells were washed three times with 3 ml of PBS and then solubilized in 0.1 M NaOH containing 0.125% Triton X-100 for protein quantification (Bradford).

#### Organization of Actin Cytoskeleton

Visualization of of actin filamnets was performed as described in [Domanski *et al.*, 2016]. Briefly, HAP1 cells, both control and *Setd3* KO clones, were cultured in IMDM supplemented with 10% fetal bovine serum, penicillin (100 U/ml) and streptomycin (100 μg/ml) at 37°C in a humidified atmosphere containing 5% CO_2_. For confocal microscopy, 5 × 10^5^ cells were seeded on a 22 × 22-mm glass coverslip in a 35-mm plate. Either twenty-four or forty-eight hours after cell seeding, culture medium was removed and cells were washed three times with PBS, fixed with 3% paraformaldehyde for 10 min in room temperature, permeabilized with 0.05% Triton X-100 in PBS for 3 minutes and washed again three times with PBS. Filamentous actin (F-actin) was visualized by staining with TRITC-phalloidin (Sigma-Aldrich) in PBS for 1 hour at room temperature. Nuclei were localized with Hoechst 33342 dye (Sigma-Aldrich). After rinsing in PBS, coverslips were mounted using fluorescence mounting medium (DakoCytomation-Agilent, Santa Clara, USA). Samples were analyzed with the use of Zeiss Axio Observer.Z1 LSM 700 confocal laser scanning microscope equipped with a Plan-Apochromat 63/1.40 Oil objective (Zeiss, Oberkochen, Germany). For detecting fluorescence of TRITC-phalloidin, 555 nm excitation line was used, and for detecting blue fluorescence of Hoechst 33342, 405 nm excitation line was employed. For 3D analysis, confocal Z-stacks comprising 10-13 optical slices were reconstructed into 3D image using Zeiss Zen 2012 software with default settings.

### Phylogenetic Analysis

Sequences homologous to rat SETD3 protein (NCBI Reference Sequence: XM_002726774.2) were identified by Protein Blast searches. A phylogenetic analysis was performed on the Phylogeny.fr platform (http://www.phylogeny.fr) [Dereeper *et al.*, 2008]. Amino acid sequences were aligned with MUSCLE (v3.7) [Edgar, 2005]. After alignment, ambiguous regions were removed with Gblocks (v0.91b) [Castresana, 2000]. Phylogenetic trees were generated using the Phylogenetic estimation using Maximum Likelihood (PhyML) [Guindon and Gascuel, 2003] with the WAG model for amino acid substitution [Whelan and Goldman, 2001] A general empirical model of protein evolution derived from multiple protein families using a maximum-likelihood approach. Mol. Biol. Evol. 18, 691-699). The final tree was customized with the editing interface TreeDyn [Chevenet *et al.*, 2006]. A confidence level was assessed using the aLRT test (minimum of SH-like and χ^2^-based parametric) [Anisimova and Gascuel, 2006].

### Analytical Methods

Protein concentration was determined spectrophotometrically according to Bradford [1976] using bovine γ-globulin as a standard. When appropriate, the His_6_-tagged recombinant proteins were detected by western blot analysis as described previously [Drozak *et al.*, 2013].

### Calculations

V_max_, K_M_ and k_cat_ for the methyltransferase activity of studied enzymes were calculated with Prism 4.0 (GraphPad Software, La Jolla, USA) using a nonlinear regression.

## RESULTS

### Purification and Identification of Rat Actin-specific Histidine N-methyltransferase

Actin-specific histidine *N*-methyltransferase activity was assayed by measuring the incorporation of [^3^H]methyl group of [^3^H]SAM into either homogeneous recombinant human β-actin produced in *E. coli* or its mutated form exhibiting a substitution of H73 by Ala residue (H73A). The H73A β-actin served as a negative control, ensuring that only activity of H73-specific *N*-methyltransferase was followed. The use of human protein as the rat methyltransferase substrate was reasonable, as the sequences of human and rat β-actin are indistinguishable due to an extreme evolutionary conservation of this protein. The enzyme was purified approximately 1200-fold from rat leg muscle by a procedure involving KCl extraction and PEG 4000 precipitation of the activity present in myofibrillar fraction of the tissue followed by its chromatography on DEAE-Sepharose, Q-Sepharose, Phenyl-Sepharose, HiScreen Blue-Sepharose, Superdex 200 and Reactive Red 120-agarose (Table 2). The methyltransferase was eluted as a single peak in each of the purification steps (Fig. 3), indicating the presence of a single enzyme species. No methylation of H73A β-actin was detected in any purification step, suggesting that the miofibrillar extract of rat muscle is devoid of other β-actin-methylating enzymes. The overall yield of the purification was only about 0.6% (cf. Table 2) due to the fact that in the last two steps of purification, only the most active fractions were used for the next step of procedure.

**Figure 3.**
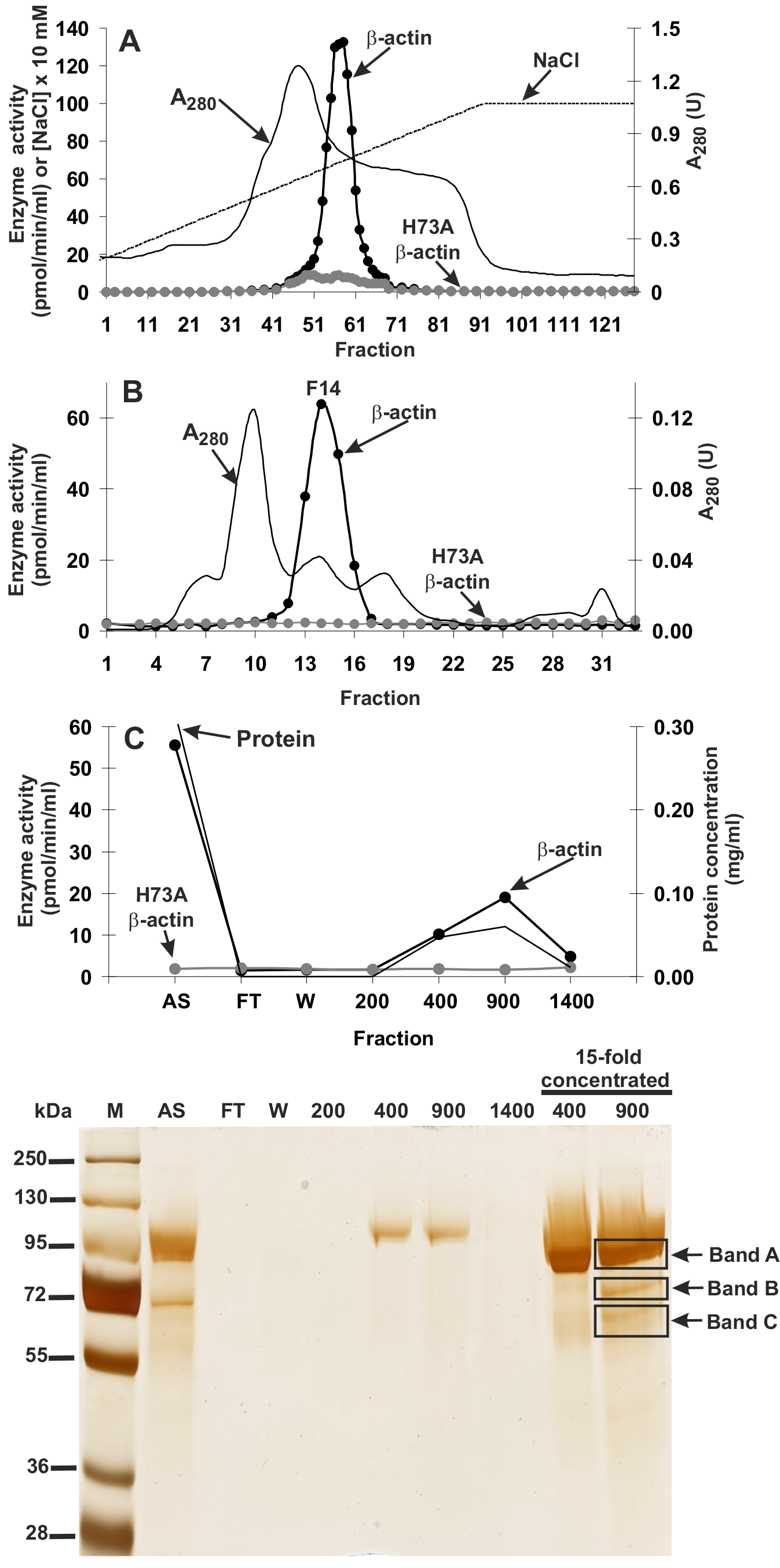
Purification of the rat actin-specific histidine *N*-methyltransferase. The enzyme was purified by chromatography on DEAE-Sepharose (A), Q-Sepharose (not shown), Phenyl-Sepharose (not shown), HiScreen Blue-Sepharose (not shown), Superdex 200 (B), and Reactive Red 120 Agarose (C) as described in “Experimental Procedures” section. Fractions were tested for the actin-specific histidine *N*-methyltransferase activity using either the homogeneous recombinant rat β-actin or its mutated form exhibiting a substitution of H73 by Ala residue (H73A). Protein concentration was determined in the most active fractions with the Bradford assay. The indicated fractions eluted from the Reactive Red 120 Agarose column were analyzed by SDS-PAGE and the gel was stained with silver [Shevchenko *et al.*, 1996]. M, prestained protein marker; AS, applied sample (Fraction 14 from Superdex 200); FT, flow through; W, wash. Fractions 200, 400, 900 and 1400 were eluted with 200, 400, 900 and 1400 mM NaCl, respectively. Additionally, fractions 400 and 900 were concentrated 15-fold by ultrafiltration (Vivaspin 500) and analyzed. The indicated bands were cut out of the gel, submitted to trypsin digestion, and analyzed by mass spectrometry.

**Table 2.** Purification of actin-specific histidine *N*-methyltransferase from rat skeletal muscles.

| Fraction | Volume | Total protein | Total activity | Specific activity | Purification | Yield |
| --- | --- | --- | --- | --- | --- | --- |
|  | *ml* | *mg* | *pmol min<sup>-1</sup>* | *pmol min<sup>-1</sup> mg<sup>-1</sup>* | *-fold* | *%* |
| Myofibrillar extract<br>(20 000 x g supernatant) | 2400 | 35273 | 8768 | 0.248 | 1 | 100 |
| 20% PEG fraction | 1000 | 16333 | 4853 | 0.297 | 1.2 | 55.4 |
| DEAE Sepharose | 84.5 | 2757 | 5753 | 2.09 | 8.4 | 65.7 |
| Q Sepharose | 63.7 | 800 | 4507 | 5.637 | 22.7 | 51.5 |
| Phenyl Sepharose | 93.4 | 66 | 964 | 14.602 | 58.8 | 11.0 |
| HiScreen Blue | 27 | 14.3 | 682 | 47.6 | 191.7 | 7.8 |
| Superdex 200# | 3 | 1.20 | 185.5 | 154.8 | 623.4 | 2.12 |
| Reactive Red 120# | 3 | 0.18 | 52.04 | 291.2 | 1172.5 | 0.59 |

<sup>#</sup> The data represent mean values for the most purified fraction of the step applied.

**Table 3.** Proteins identified in the gel bands submitted to trypsin digestion and MS/MS analysis. Identified proteins are listed for each band according to their score calculated by ProteinLynx Global Server software (PLGS). For each protein, its molecular weight (mW) and the sequence coverage are also indicated. Occasional peptide hits corresponding to keratins have not been included.

| **Gel band** | **Protein name** | **NCBI Protein accession number** | **PLGS score<sup>a</sup>** | **mW (Da)** | **Coverage (%)** |
| --- | --- | --- | --- | --- | --- |
| **A** | Elongation factor 2 | NP_058941.1 | 16249 | 95222 | 73.5 |
| **B** | Calpain 1 catalytic subunit | NP_062025.1 | 3024 | 82067 | 47.7 |
|  | Elongation factor 2 | NP_058941.1 | 789 | 95222 | 15.2 |
|  | Nuclear protein localization protein 4 homolog | NP_542144.1 | 778 | 68013 | 31.7 |
| **C** | Elongation factor 2 | NP_058941.1 | 2024 | 95222 | 42.1 |
|  | Nuclear protein localization protein 4 homolog | NP_542144.1 | 1263 | 68013 | 37.3 |
|  | Histone lysine N methyltransferase setd3 | XP_002726820.2 | 1136 | 67378 | 39.1 |
|  | Eukaryotic peptide chain release factor GTP binding subunit ERF3A | NP_001003978.1 | 505 | 68708 | 41.2 |
|  | Guanylate binding protein 1 | NP_598308.1 | 404 | 67312 | 21.8 |
|  | Filamin B like isoform X2 | XP_003751493.2 | 272 | 121901 | 1.4 |

<sup>a</sup>PLGS Score is calculated by the ProteinLynx Global Server (v2.4) software using a Monte Carlo algorithm to analyze all acquired mass spectral data and it is a statistical measure of accuracy of assignment. A higher score implies a greater confidence of protein identity.

SDS-PAGE analysis of the peak activity fractions (400 and 900) derived from the Reactive Red 120-agarose revealed a low protein content resulting in very faint protein bands, which prevented the identification of protein bands coeluting with the enzyme activity (cf. Fig. 3). Thus, peak activity fractions 400 and 900 were concentrated about 15-fold and reanalyzed together with non-concentrated ones by SDS-PAGE. Actin-methylating activity was coeluted with three protein bands of about 65, 75 and 90 kDa, well-visible in the concentrated fraction 900 (cf. Fig. 3). The bands were cut out from the gel and digested with trypsin. The resulted peptides were analyzed by MS/MS and compared with the reference proteome of rat obtained from NCBI Protein database. The analysis indicated that only band “C” contained a methyltransferase designated as histone lysine *N*-methyltransferase SETD3 (Table 2). Eighteen matching peptides (underlined in Fig. 4) were found to cover about 39% of its sequence. To exclude the possibility of missing any potential methyltransferase due to a poor extraction of tryptic peptides from protein bands, we performed MS/MS identification of all proteins present in the entire fraction 14 (Superdex 200) and fraction 900 (Reactive red 120), and SETD3 protein was again the only methyltransferase found in these fractions (not shown).

**Figure 4.**
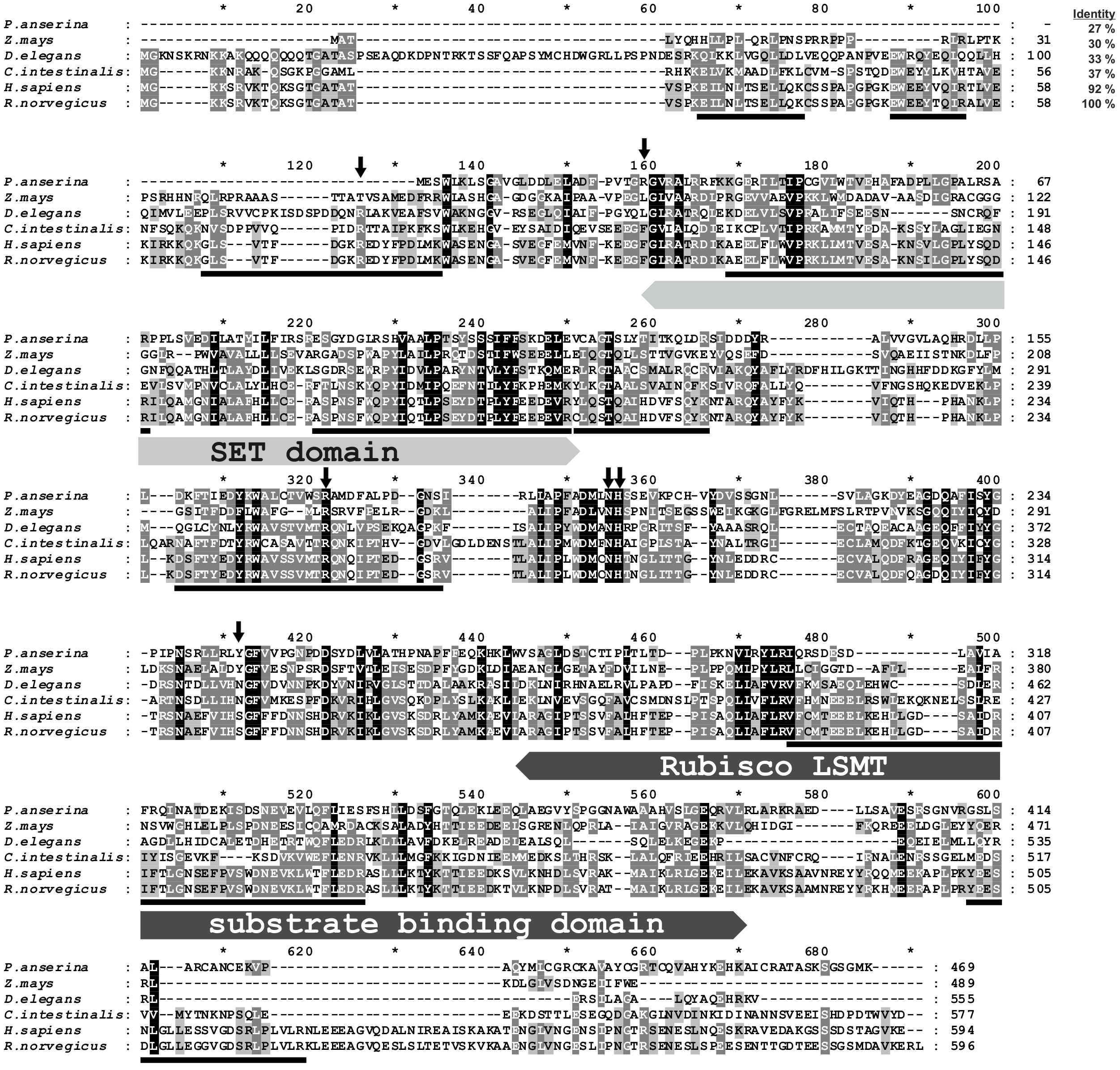
Amino acid sequence alignment of the rat SETD3 protein with its orthologs. Sequences of rat (*R. norvegicus*, XP_002726820.2), human (*H. sapiens*, NP_115609.2), *Ciona intestinalis* (*C. intestinalis*, XP_002131202.1), *Drosophila elegans* (*D. elegans*, XP_017114801.1), *Podospora anserina* (*P. anserina*, CDP29262.1) and *Zea mays* (*Z. mays*, NP_001168589.1) protein were obtained from NCBI Protein database. Both rat and human sequences have been confirmed by PCR amplification of the cDNA and DNA sequencing. Percentage of amino acid identities with the rat SETD3 protein is given in the upper right. The conserved protein substrate-binding domains (SET and Rubisco LSMT substrate binding) are labeled above the alignment, while amino acid residues interacting with *S*-adenosyl-L-methionine (SAM) are indicated by arrows as inferred form the crystal structure of human SETD3 enzyme (PDB 3SMT). The peptides identified by mass spectrometry in the protein purified from rat leg muscle are underlined in the rat sequence. Level of residues conservation is indicated by black (100%), dark grey (70% and more) and light gray (50% and more) background.

As SETD3 protein was shown to methylate histones [Eom *et al.*, 2011] and all SET-domain containing enzymes characterized to date are protein lysine methyltransferases [Clarke, 2013], it seemed very unlikely that SETD3 is protein histidine *N*-methyltransferase. We therefore performed another complete, though modified, round of purification of the actin-methylating enzyme from rat leg muscles, and MS/MS analysis of the most purified fractions from Superdex 200 yielded again SETD3 protein as the only meaningful candidate for rat actin-specific histidine *N*-methyltransferase (results not shown). This finding was in agreement with previous reports, suggesting that actin-specific histidine *N*-methyltransferase corresponded to ≈70-kDa protein [Raghavan, 1992].

### Analysis of SETD3 Protein Sequences

Protein BLAST searches [Altschu *et al.*, 1997] with rat SETD3 and phylogenetic analysis of resulting sequences indicated that orthologs of this protein were found in almost all eukaryotes including vertebrates (≈75-98% identity with the rat sequence), insects (≈35-40% identity), plants and fungi (≈30% identity) (Fig. 5). No SETD3 enzymes was detected in the proteome of *Saccharomyces cerevisiae* and of *Naegleria gruberi* in agreement with previous reports, showing the absence of actin-specific histidine methyltransferase activity in these species [Kalhor *et al.*, 1999]. All taxa followed the expected lines of descent, indicating that the enzyme was already present in a common eukaryotic ancestor (cf. Fig. 5). A low similarity in amino acid sequence between eukaryotic SETD3 orthologs and some bacterial proteins was also detected (e.g. ≈30% identity, *Cystobacter fuscus*, NCBI Reference Sequence: WP_095987699.1), suggesting that the eukaryotic enzyme might have been acquired from an ancestral prokaryote.

**Figure 5.**
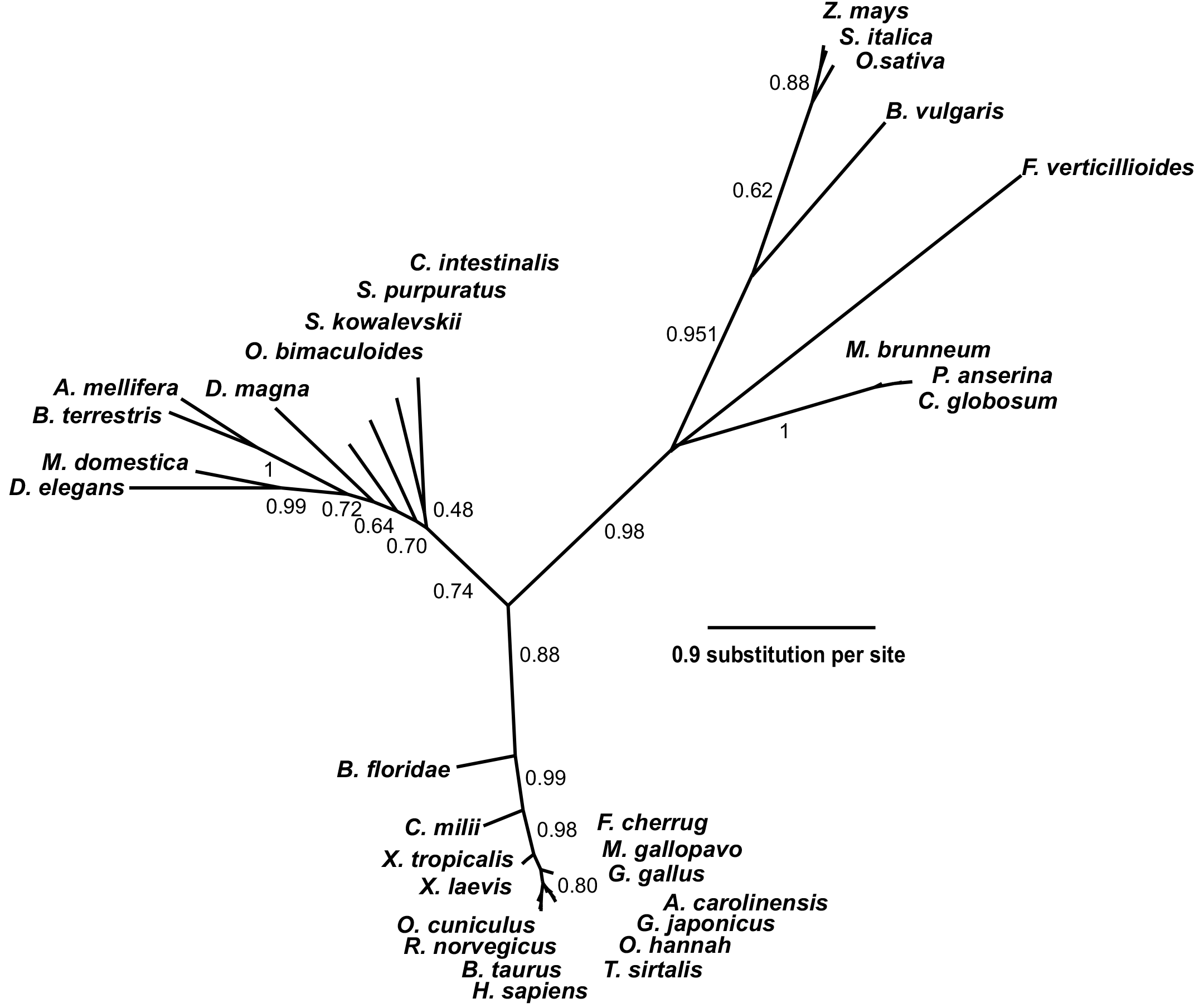
Phylogenetic tree of the SETD3 proteins. Protein sequences were aligned using Muscle [Edgar, 2004], while the phylogenetic tree was inferred with the use of PhyML [Guindon and Gascuel, 2003] implemented in phylogeny.fr web service [Dereeper *et al.*, 2008]. Brunch support values assessed using the aLRT test are indicated [Anisimova and Gascuel, 2006]. The protein sequences used for the analysis are as follows: *Anolis carolinensis* (XP_003214383.2); *Apis mellifera* (XP_016770011.1); *Beta vulgaris* (XP_010683122.1); *Bombus terrestris* (XP_003398458.1); *Bos taurus* (XP_589822.3); *Branchiostoma floridae* (XP_002596839.1); *Callorhinchus milii* (XP_007906724.1); *Chaetomium globosum* (XP_001224775.1); *Ciona intestinalis* (XP_002131202.1); *Daphnia magna* (KZS12928.1); *Drosophila elegans* (XP_017114801.1); *Falco cherrug* (XP_005438913.1); *Fusarium verticillioides* (XP_018752240.1); *Gallus gallus* (NP_001006486.1); *Gekko japonicus* (XP_015275964.1); *Homo sapiens* (NP_115609.2); *Meleagris gallopavo* (XP_003206761.2); *Metarhizium brunneum* (XP_014542924.1); *Musca domestica* (XP_011294563.1); *Octopus bimaculoides* (XP_014787293.1); *Ophiophagus hannah* (ETE71402.1); *Oryctolagus cuniculus* (XP_008247172.2); *Oryza sativa* (XP_015651332.1); *Podospora anserina* (CDP29262.1); *Rattus norvegicus* (XP_002726820.2); *Saccoglossus kowalevskii* (XP_006819296.1); *Setaria italica* (XP_004956796.1); *Strongylocentrotus purpuratus* (XP_798530.2); *Thamnophis sirtalis* (XP_013914404.1); *Xenopus laevis* (OCT68299.1); *Xenopus tropicalis* (XP_012823880.1); *Zea mays* (NP_001168589.1).

All identified proteins contain the SET and Rubisco LSMT substrate-binding domains at their N- and C-terminus, respectively. The SET domain is thought to interact directly with protein substrate and most likely contributes indirectly to SAM binding as well, while Rubisco LSMT substrate-binding domain seems to interact with a protein substrate only (*cf.* Figs. 4 and 6). Although the amino acid sequence identity between orthologs from distinct species such as rat and maize (*Zea mays*) was rather low (about 30%), the alignment of their sequences showed the presence of several motifs that were strictly or at least moderately conserved in both domains, suggesting that these orthologs may have similar sort of substrates (*cf.* Fig. 4).

Partial protein structure of human SETD3 has been published (PDB 3SMT). It lacks about one hundred amino acids at C-terminus, though both substrate binding domains are well-preserved. As shown in Fig. 6, these two domains form a wide cleft in the center of SETD3, making that the enzyme adopts a cradle-like shape. The SAM molecule is bound in the bottom of the cleft, exposing its labile methyl group (*cf.* Fig. 6). It seems likely that the cradle-shape of the enzyme allows it to arch over a protein substrate, and the position of SAM facilitates the subsequent transfer of the methyl group onto a specific amino acid residue, most likely located at the surface of a substrate protein.

**Figure 6.**
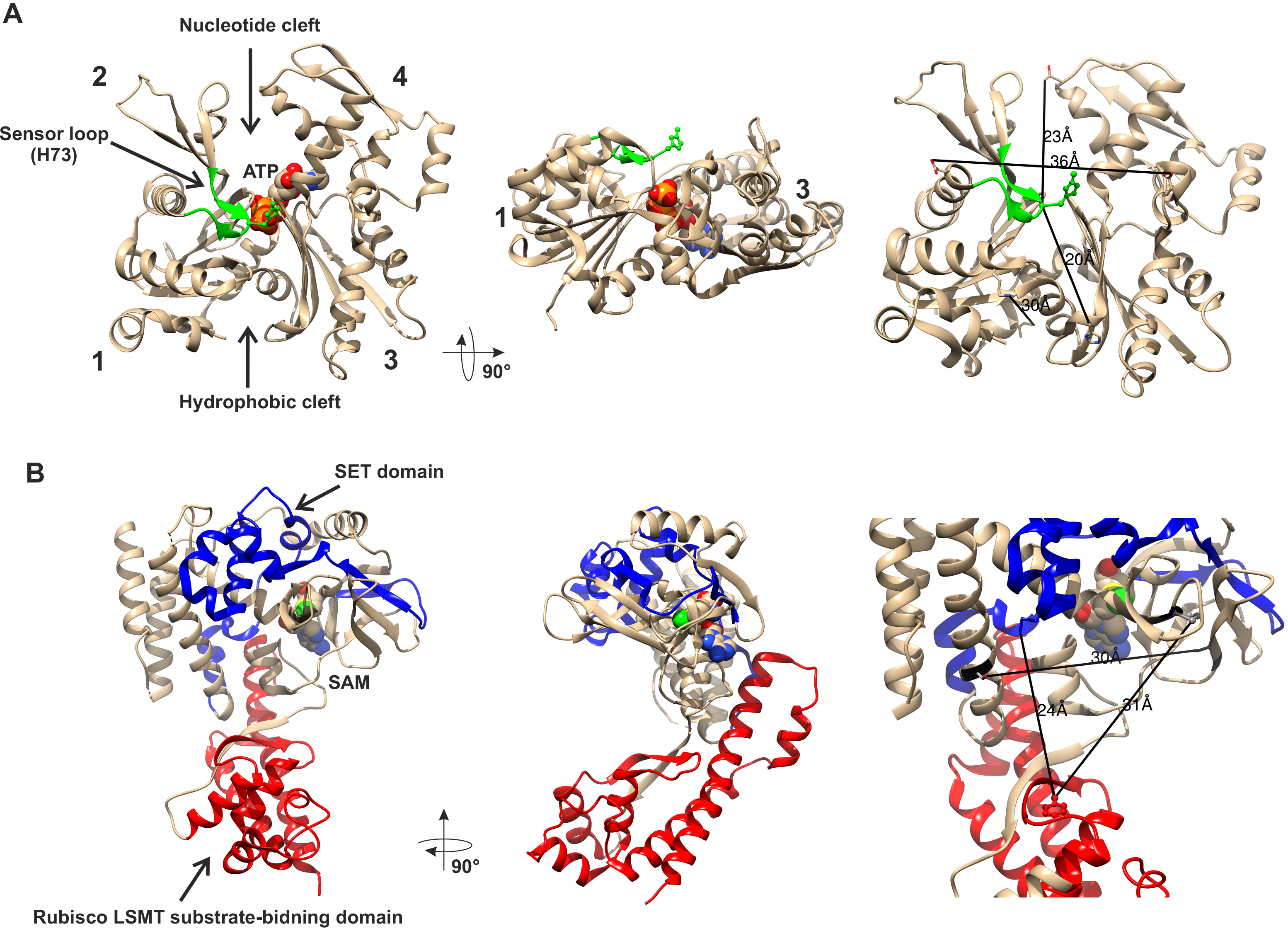
Structures of human β-actin and SETD3 enzyme. Ribbon representations of the structures of both the actin monomer (A) and the SETD3 protein (B) are shown in different projections. (A) The four subdomains of actin are marked with numbers. Subdomains 1 and 2 form the small (outer) domain, whereas subdomains 3 and 4 constitute the large (inner) domain. Two large clefts are present between these domains: the nucleotide binding cleft and the hydrophobic one, while ATP is bound at the center of the molecule. The nucleotide sensor loop that contains the methyl-accepting H73 residue, spans from Pro70 to Asn78 and is shown in green. (B) The SET and Rubisco LSMT substrate-binding domains of SETD3 are highlighted in blue and red respectively. The spatial organization of these domains gives the enzyme a cradle-like shape. The *S*-adenosyl-L-methionine (SAM) molecule is bound at the bottom of this cradle, exposing its labile methyl group (green) to a protein substrate. Several residue-residue distances are shown to give information concerning the molecular dimension of these proteins. All models were prepared using UCSF Chimera [Pettersen *et al.*, 2004] from the Protein Data Bank (PDB) structures of β-actin (2BTF) and SETD3 (3SMT).

As shown in Fig. 6, β-actin has a globular shape, flattened in one dimension. It consists of large and small domains, when considering their size. The small and large domains are further subdivided into subdomains 1, 2 and 3, 4, respectively. The nucleotide binding cleft (ATP or ADP) is located inside the cleft between subdomains 2 and 4, while a hydrophobic cleft is present between subdomains 1 and 3 on the “bottom” of the actin molecule, and serves as a primary binding site for numerous actin-binding proteins. The methyl-accepting H73 residue is present in a nucleotide sensor loop (Pro70 to Asn 78) that forms an insert between subdomains 1 and 2. The His73 is present on the surface of actin monomer and seems to be easily accessible for the methyltransferase. Furthermore, roughly estimated dimensions of actin monomer and the substrate biding cleft of the SETD3 enzyme indicates that actin should accommodate well at the cleft of the methyltransferase and steric hindrance should not prevent interactions between these proteins.

### Characterization of Recombinant SETD3 Proteins

To confirm the molecular identity of rat actin-specific histidine *N*-methyltransferase with that of rat SETD3 protein and to compare the enzymatic activities of rat and human methyltransferase, these two SETD3 orthologs were expressed in either HEK293T and COS-7 cells as fusion proteins with the C-terminal polyhistidine tag (Fig. 7). The recombinant enzymes catalyzed methylation of recombinant β-actin, and not its mutated form H73A, as determined by the radiochemical assay (cf. Fig. 7). For further studies, large amounts of recombinant enzymes were produced in COS-7 cells, purified to homogeneity (*cf.* Fig. 1) and shown to catalyze the modification of actin (Fig. 8).

**Figure 7.**
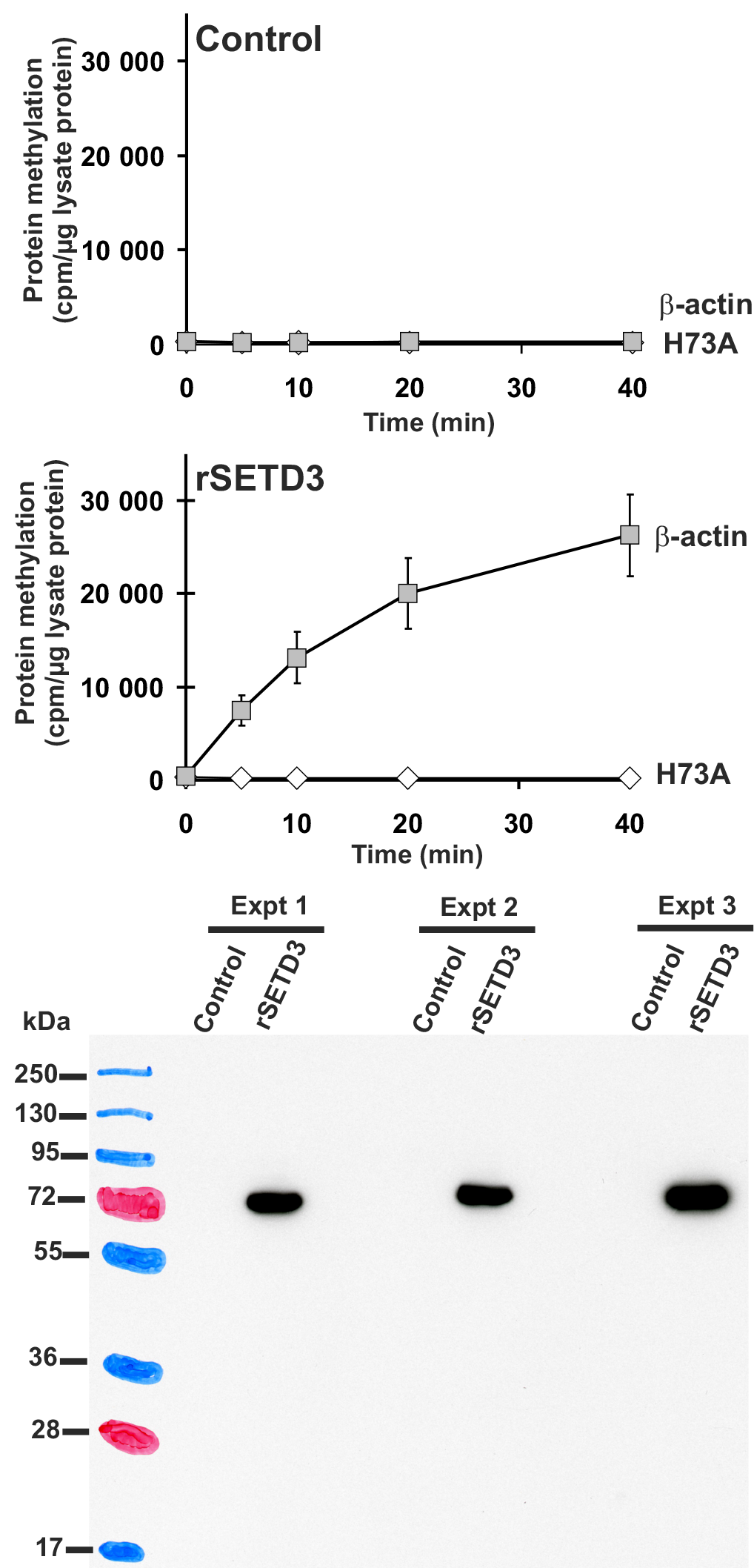
Time course of β-actin methylation in lysates of the control and the SETD3-overexpressing COS-7 cells. COS-7 cells were transfected for 48 h with an empty vector (Control) or a vector encoding the rat SETD3 protein (rSETD3). Cell-free lysates (3 μg of protein) were incubated at 37 °C for the indicated times in the presence of 1 μM [^1^H+^3^H]SAM (100 pmol, ≈450 × 10^3^ cpm) and 2 μM (200 pmol, 8.9 μg) homogenous recombinant rat β-actin or its mutated form in which H73 was replaced by alanine residue (H73A). Proteins were precipitated with 10% trichloroacetic acid to determine the incorporation of radioactivity. The presence of recombinant protein in tested lysates was verified by Western blot analysis, which was performed using 20 μg of total protein and an antibody against the His_6_ tag. Values are the means ± S.E. (error bars) of 3 independent transfections. When no error bar is shown, the error is smaller than the width of the line.

**Figure 8.**
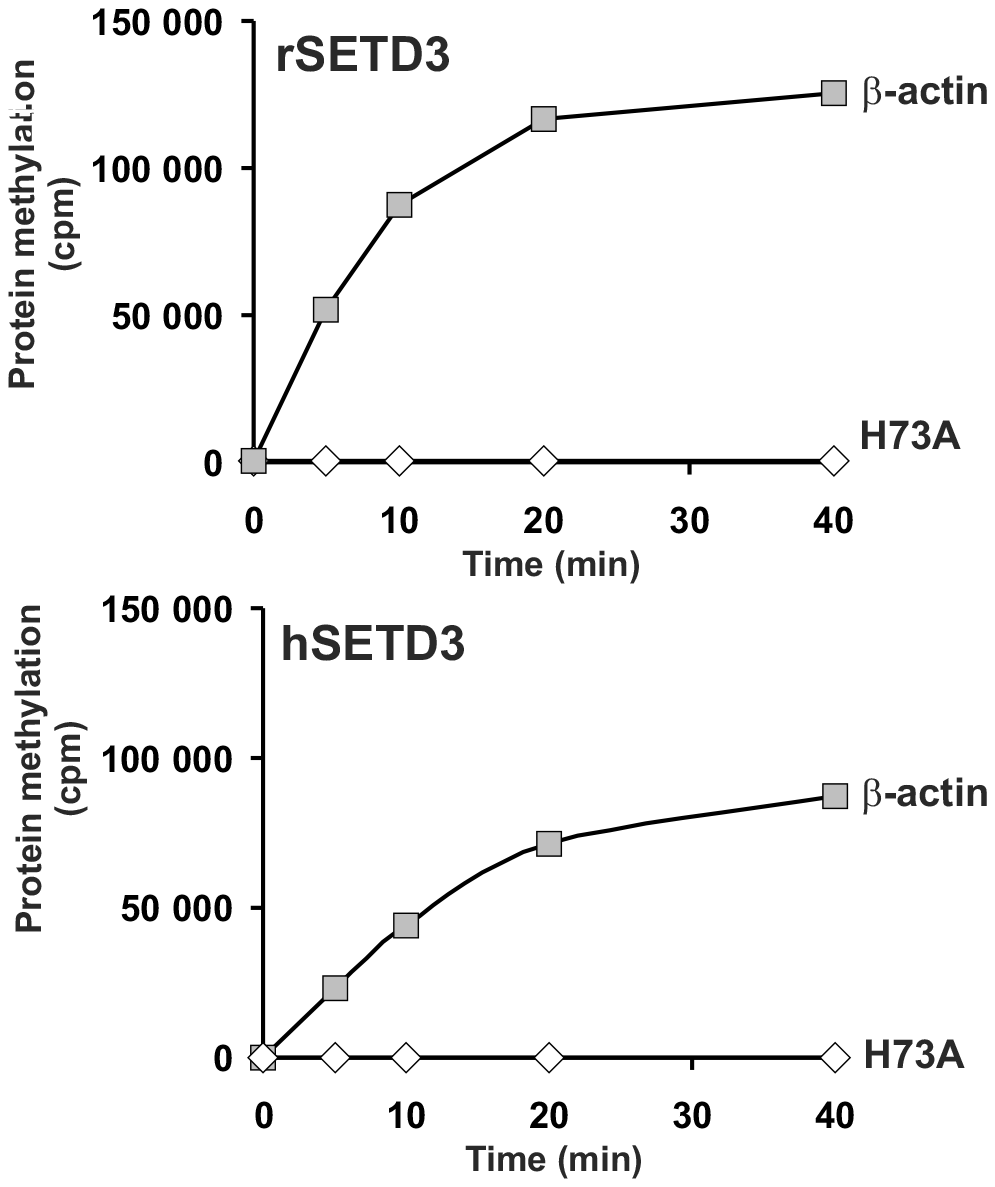
Time course of β-actin methylation in the presence of purified recombinant rat or human SETD3 enzyme. Homogeneous recombinant rat SETD3 (rSETD3, 0.4 μg protein) or its human orthologue (hSETD3, 0.3 μg protein) were incubated at 37 °C for the indicated times in the presence of 1 μM [^1^H+^3^H]SAM (100 pmol, ≈300 - 400 × 10^3^ cpm) and 2 μM (200 pmol, 8.9 μg) homogenous recombinant rat β-actin or its mutated form in which H73 was replaced by alanine residue (H73A). Proteins were precipitated with 10% trichloroacetic acid to determine the incorporation of radioactivity. The figure shows results of representative experiments.

Taking into account that SETD3 actively methylates recombinant β-actin, we tested whether the enzyme can modify a synthetic peptide, corresponding to the H73-containg nucleotide sensor loop of actin molecule (Peptide H: YPIE**H**GIVT) as it had been previously shown for a partially purified rabbit actin-specific histidine *N*-methyltransferase [Raghaven *et al.*, 1992]. As presented in Fig. 9, the two SETD3 orthologs catalyzed methylation of peptide H, while they were inactive towards peptide A (YPIE**A**GIVT) - a variant of the nucleotide sensor loop, exhibiting substitution of H73 by alanine residue. Though their activities towards peptide H were very poor (≈ 0.02 nmol × min^-1^ × mg^-1^ enzyme protein) in comparison with the protein substrate (≈ 5 nmol × min^-1^ × mg^-1^ enzyme protein), these findings provided further indication that SETD3 is the histidine-specific *N*-methyltransferase, since histidine is the only amino acid residue present in the peptide H that could potentially accept methyl group.

**Figure 9.**
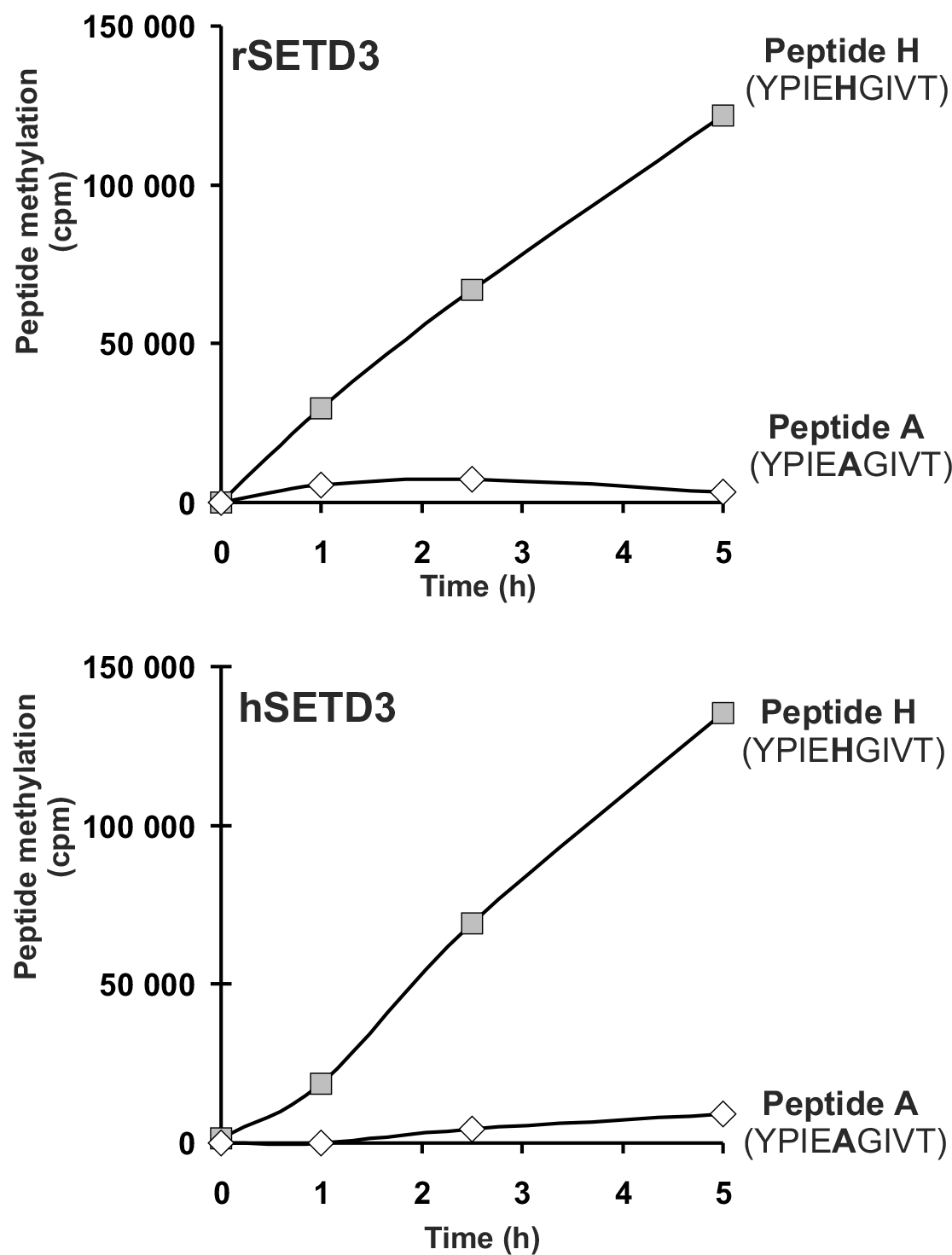
SETD3-dependent methylation of synthetic peptides - analogues of the methylation site in β-actin. Homogeneous recombinant rat SETD3 (rSETD3, 5.8 μg protein) or its human orthologue (hSETD3, 4.6 μg protein) were incubated at 37 °C for the indicated times in the presence of 1 μM [^1^H+^3^H]SAM (100 pmol, ≈350 − 400 × 10^3^ cpm) and 2 mM (200 nmol) synthetic peptides corresponding to residues 69-77 of either β-actin (Peptide H: YPIE*H*GIVT) or its mutated form (Peptide A: YPIE**A**GIVT). Methylated peptides were separated from [^1^H+^3^H]SAM employing the sample fractionation on Dowex 50W columns. The figure shows results of representative experiments.

To confirm the specific methylation of H73 of the recombinant human β-actin, homogenous recombinant rat and human SETD3 enzymes were incubated with β-actin in the presence of deuterated SAM ([^2^H]SAM), yielding the extent of actin methylation equal to about 35%, as determined in parallel radiochemical assay. The protein products of the reaction were separated by SDS-PAGE, and bands corresponding to the β-actin polipeptide were cut out from the gel and digested with trypsin. Resulted peptides were analyzed by nanoUPLC-Q-TOF and compared with a protein databank supplemented with amino acid sequence of the recombinant β-actin. Peptide YPIEHGIVTNWDDMEK (M+H = 1963.9355 Da) comprising His73 residue was the only tri-deuterium-methylated peptide detected by PLGS software. The manual inspection of MS/MS spectra of both the parent ion of this peptide and its fragments confirmed the presence of tri-deuterium-methylated His73 (Fig. 10). Both rat and human SETD3 enzymes catalyzed tri-deuterium-methylation of His73 in β-actin, while no such modification was detected in YPIEHGIVTNWDDMEK peptide (M+H = 1946.9021 Da) or any other tryptic peptide derived form β-actin protein incubated with no enzymes in the reaction mixture (*cf.* Fig. 10).

**Figure 10.**
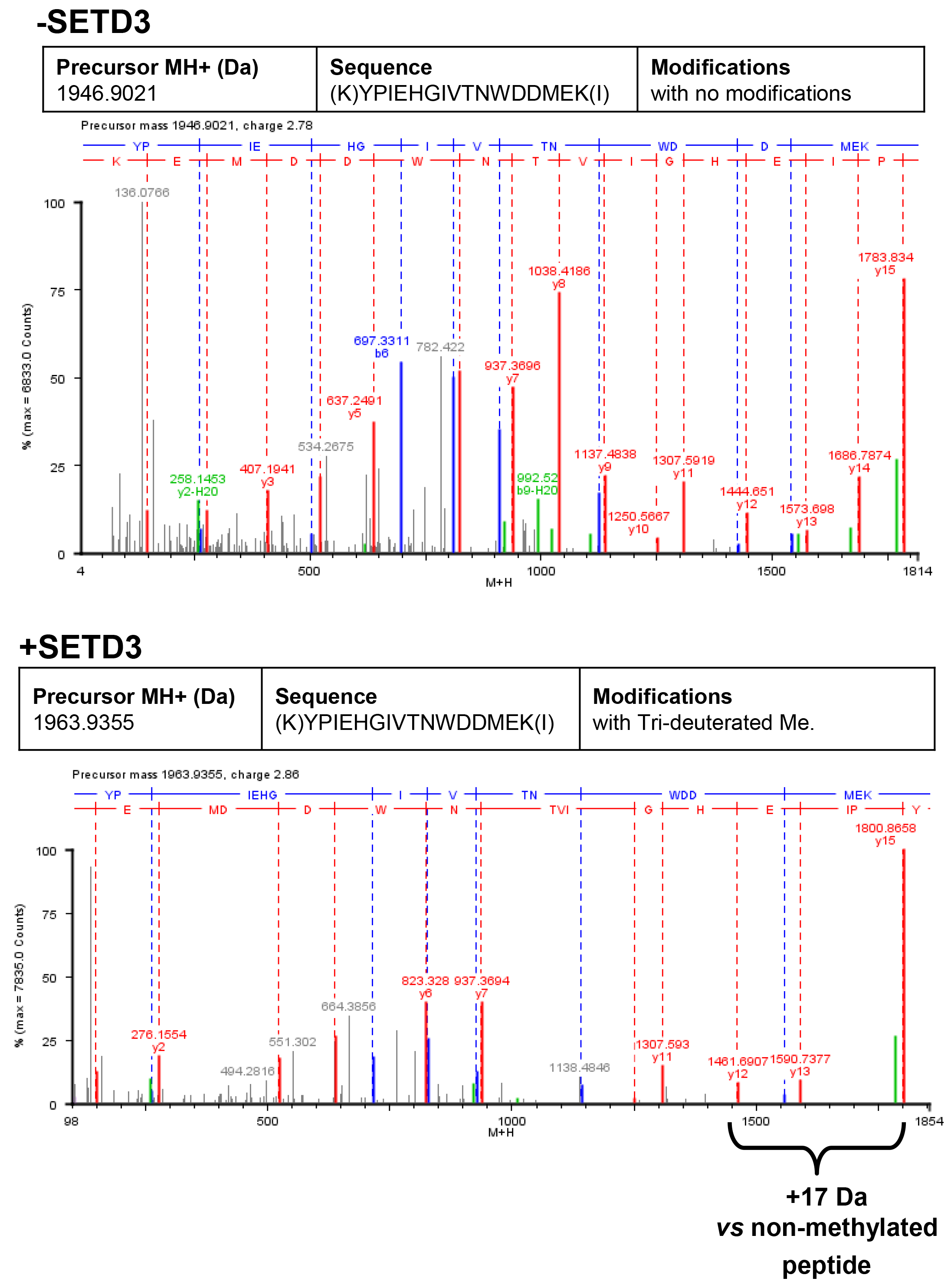
Deconvoluted Q-TOF spectra of both non-modified and deuterium(D_3_)-methylated β-actin peptides produced by the human SETD3 protein. Homogenous recombinant human β-actin (6.4 μg) was incubated for 90 min at 37°C in a reaction mixture containing 20-30 μM ([^2^H]SAM), 4.5-9.0 μg SAH nucleosidase and 1.75-3.5 μg adenine deaminase in either the absence (−SETD3) or presence of 1.1-3.5 μg the homogenous recombinant human SETD3 protein (+SETD3). Following the reaction, β-actin was separated from other proteins by SDS-PAGE and analyzed by tandem mass spectrometry as described in “Experimental Procedures”. Detection of the trideuterium-methylated peptides was performed in a fully automatic mode by ProteinLynx Global Server 2.4 software (Waters, USA), and the trideuterium-methylation (+17,03448 Da) of Cys, Asp, Asn, His, Lys, Arg, Glu, Gln residues of tryptic peptides was examined. None of the (trideuterium−)methylated peptides was detected in the control reaction (−SETD3), whereas the H73-comprising peptide: YPIEHGIVTNWDDMEK (M+H = 1946.9021 Da, y16 ion) was the only trideuterium-methylated peptide (M+H = 1963.9355 Da, y16 ion) in the presence of SETD3. The mass shift (+17.03448 Da) was detected only in His-containing peptide fragments (y12, y13, y15, y16), indicating His73 residue as the site of methylation. The figure shows results of the representative experiment. Two independent methyl(D_3_)-labeling reactions were performed.

The kinetic properties of homogenous recombinant SETD3 proteins were studied with the homogenous recombinant human β-actin in the presence of *E. coli* SAH nucleosidase and *B. subtilis* adenine deaminase. The bacterial enzymes served to break down S-adenosyl-L-homocysteine (SAH), a potent inhibitor of methyltransferases, preventing from its accumulation in the reaction mixture [Dorgan *et al.*, 2006]. Both auxiliary enzymes were added in ≈100-fold molar excess in comparison to SETD3 enzymes. As shown in Table 4, rat and human enzymes exhibited very high affinity towards both SAM and the protein substrate, presenting K_M_ values much lower than the physiological concentrations of SAM and β-actin monomers in vertebrate cells (≈ 30 μM and 50-100 μM, respectively [Clarke and Banfield, 2001; Pollard, 2017]). These two enzymes displayed comparable V_max_ values estimated at saturating SAM concentrations, while the human SETD3 showed about 4-fold lower K_M_ for β-actin (≈0.8 μM) than the rat enzyme (3 μM). The catalytic efficiency (K_M_/k_cat_) to β-actin for human SETD3 was therefore 3-fold higher than the value determined for the rat enzyme (cf. Table 4). However, taking into account the intracellular concentrations of SAM and β-actin monomers, the two enzymes operate probably at similar V_0_ *in vivo*, i.e. close to their V_max_ (≈10 nmol × min^-1^ × mg^-1^ protein).

**Table 4.** Analysis of β-actin methylation at His73 residue *in vivo* by tandem mass spectrometry. The total number of identified peptide sequences with His73 (peptide spectrum matches, PSMs) for β-actin isolated from either Wild-type HAP1 cells (WT-A2, WT-A6) or *Setd3*-deficient clonal HAP1 cells (KO-A1, KO-A3) are indicated. The total number of PSMs recorded for β-actin (Uniprot, P60709) in these MS/MS experiments is as follows: 301 (WT-A2), 387 (WT-A6), 485 (KO-A1), 529 (KO-A3).

| Peptide variant | Number of identified peptide sequences (PSMs) |  |  |  |
| --- | --- | --- | --- | --- |
|  | WT-A2 | WT-A6 | KO-A1 | KO-A3 |
| His73 | ND | 2 | 30 | 30 |
| Methylated His73 | 20 | 19 | 3 | 4 |

ND, not detected.

It is well-established that actin produced in bacteria forms non-native misfolded species that aggregate into inclusion bodies [Stemp *et al.*, 2005]. This is due to the fact that actin protein requires the assistance of the eukaryotic chaperonin CCT (TRiC) to fold to its native confirmation [Martín-Benito *et al.*, 2002; Stemp *et al.*, 2005]. However, the completely denatured actin was capable of spontaneous acquiring a large portion of its native tertiary structure [Martín-Benito *et al.*, 2002], while the refolded recombinant β-actin used in experiments described above was mostly like that in nature. As these observations raised the question about the activity of SETD3 towards the natively folded substrate, the recombinant human β-actin was produced in *S. cerevisiae* – an eukaryotic species devoid of actin-specific histidine methyltransferase activity [Kalhor *et al.*, 1999]. Unexpectedly, both rat and human SETD3 catalyzed methylation of neither ATP-β-actin nor ADP-β-actin, as determined by the radiochemical assay (< 600 cpm × 20 min^-1^ in the presence of 0.3-0.4 μg SETD3 protein, 200 pmol ATP/ADP-β-actin, 100 pmol [^1^H+^3^H] SAM, 3200-4200 cpm × pmol^-1^ SAM, *cf.* Fig. 8), suggesting that native nucleotide-bound actin monomers may not necessarily be the endogenous substrates of SETD3 methyltransferase. To throw more light on this issue, nucleotide-free actin monomers in complexes with either profilin or cofilin were prepared and tested as potential substrates. The methylation of profilin-β-actin was negligible, while cofilin-β-actin complex appeared a substrate for SETD3 enzyme, albeit very poor one (only ≈3-4% of total protein of the complex could be methylated). These findings suggest that G-actin coupled to a certain actin-bidning protein or even F-actin may be substrates for the SETD3 enzymes *in vivo*.

### Characterization of Setd3-deficient Model Human Cells

To verify the importance of SETD3 activity for the methylation of β-actin H73 residue *in vivo*, human cell lines (HAP1) deficient in SETD3 were prepared employing CRISPR/Cas9 technology. Five distinct *Setd3-KO* clones were generated, as described in *Experimental Procedures*, and three *Setd3*-deficint cell lines were ultimately selected for further studies (KO-A1, KO-A3 and KO-A5). Sequencing of the DNA indicated the presence of mutations, preventing from the production of an active SETD3 enzyme in these clones (not shown), while the absence of the enzymatic protein in *Setd3*-KO cell lysates was further confirmed by western-blot analysis (Fig. 11).

**Figure 11.**
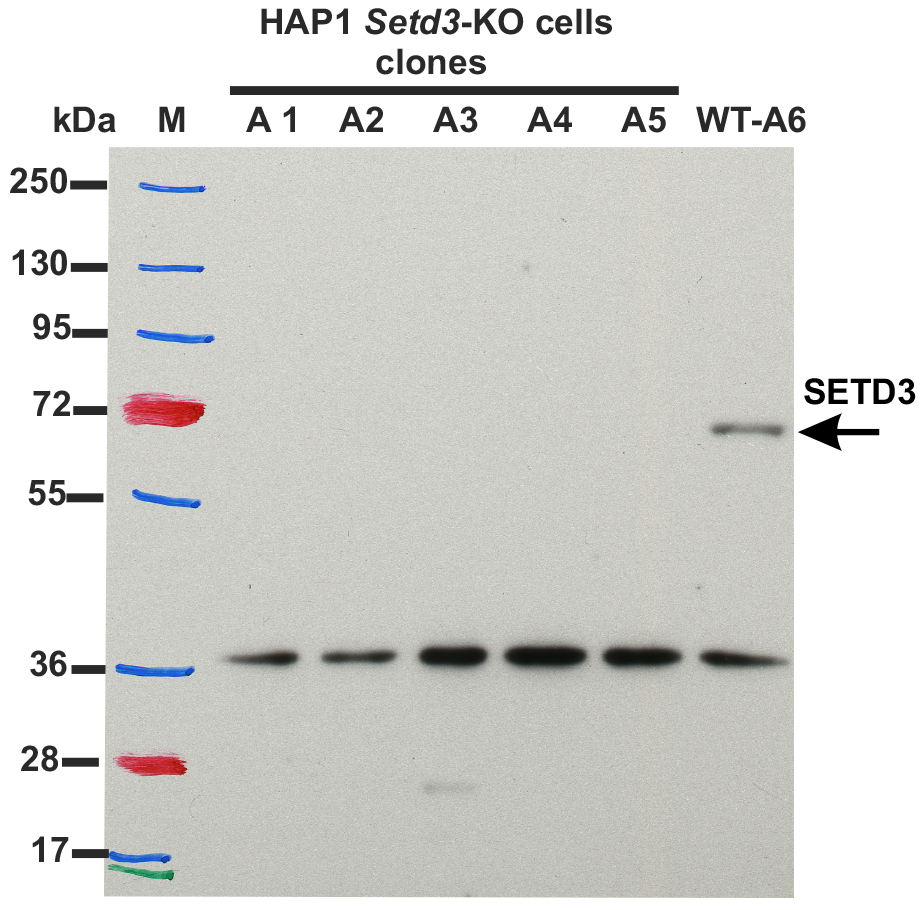
Western blot analysis of the endogenous SETD3 protein expression in control and *Setd3*-deficient HAP1 cells. HAP1 cells were transfected with a pair of plasmids pSpCas9n(BB)-2A-Puro coding for Cas9n nickase and sgRNA sequences targeting two sequences localized at exon 4 of human *Setd3* gene. After selection with puromycin, five clonal cell lines were isolated and tested for production of the SETD3 protein (≈ 70 kDa). Control HAP1 cell line (WT-A6) that has been also submitted to the CRISPR/Cas9 gene inactivation procedure, but yielding no modification of the *Setd3* gene; this was done to exclude off-target effects of the procedure on HAP1 phenotype. The analysis was performed using 50 μg of the cell lysate protein with a rabbit primary antibody against the human SETD3 (Abcam, ab174662) along with the horseradish peroxidase-conjugated goat anti-rabbit secondary antibody. The presence of nonspecific signal (≈ 36 kDa) is in agreement with the specification of primary antibody.

To determine the methylation status of β-actin H73 residue in both control and *Setd3*-deficient cells, actin was partially purified from these cell lines and analyzed by SDS-PAGE. The actin band was identified, cut out of the gel and digested with trypsin. Resulted peptides were then analyzed by tandem mass spectrometry, as described in *Experimental Procedures*. As shown in Table 5, methylation of H73 residue was detected in almost all H73-containing β-actin peptides derived from the wild-type Hap1 cells, while nearly 90% β-actin peptides derived from *Setd3*-deficient cells exhibited non-methylated H73 residue. Importantly, the total number of β-actin peptides identified in MS/MS experiments was roughly comparable for control (300-400 Peptide Spectrum Matches) and KO cells (≈500 PSM), indicating that similar amounts of actin protein were analyzed in all experiments. These data clearly show that actin-specific histidine *N*-methyltransferase activity strictly depends on the presence of SETD3 protein in HAP1 cells.

**Table 5.** Kinetic properties of rat and human SETD3 proteins. Kinetic properties were determined with the use of purified recombinant C-terminal His_6_-tagged SETD3 protein.

| Substrate | Rat SETD3 |  |  |  | Human SETD3 |  |  |  |
| --- | --- | --- | --- | --- | --- | --- | --- | --- |
| | $V_{\max}$<br>$nmol\ min^{-1}\ mg^{-1}$ | $K_M$<br>$\mu M$ | $k_{cat}$<br>$min^{-1}$ | $k_{cat}/K_M$<br>$min^{-1}\ \mu M^{-1}$ | $V_{\max}$<br>$nmol\ min^{-1}\ mg^{-1}$ | $K_M$<br>$\mu M$ | $k_{cat}$<br>$min^{-1}$ | $k_{cat}/K_M$<br>$min^{-1}\ \mu M^{-1}$ |
| β-actin | 11.280± 1.018 | 2.996 ± 0.507 | 0.80 | 0.27 | 9.091 ± 0.308 | 0.752 ± 0.070 | 0.65 | 0.86 |
| *S*-adenosyl-L- methionine | 8.053 ± 0.136 | 0.109 ± 0.008 | 0.57 | 5.23 | 8.649 ± 0.119 | 0.116 ± 0.007 | 0.61 | 5.25 |

Determinations for *S*-adenosyl-L-methionine (SAM) were performed with the SETD3 preparations (0.04-0.05 μg protein, 5-7 nM) that were incubated for 8 min at 37°C in the reaction mixture containing 5 μM recombinant β-actin and variable concentrations of [<sup>1</sup>H+<sup>3</sup>H] SAM ( $\approx 320 \times 10^3$ cpm), while the measurements for β-actin were done following a 5-min incubation of the SETD3 in the presence of 1 μM concentration of [<sup>1</sup>H+<sup>3</sup>H] SAM (100 pmol, $\approx 300 \times 10^3$ cpm). In all experiments reaction mixture contained the homogenous recombinant SAH nucleosidase (1.6 μg protein, 600 nM, *E. coli*) and adenine deaminase (3.9 μg protein, 600 nM, *B. subtilis*) to prevent *S*-adenosyl-L-homocysteine (SAH) accumulation. Values are the means of three separate experiments. The S.E. values are also given.

Methylation of actin H73 residue was shown to stabilizes actin filaments, while the absence of such modification resulted in an increased rate of filaments depolymerization *in vitro* [Nyman *et al.*, 2002]. To determine the effect of *setd3*-deficiency on the organization of actin filaments, confocal microscopy was employed to analyze control HAP1 cells and SETD3-KO cell lines, following F-actin staining with TRITC-phalloidin. To obtain a comprehensive picture of actin filaments, the morphology of F-actin was reconstructed in 3D with the use of Zeiss Zen software. As shown in Fig. 12, a clear decrease in F-actin content was visible in cells lacking SETD3 activity, suggesting an accelerated actin depolymerization and a loss of cytoskeleton integrity in SETD3-KO cells. These effects were more pronounced in cells cultured for 48h in comparison with those grown for 24h only, and seemed to be in agreement with results of previous studies performed with the use of purified recombinant actin [Nyman *et al.*, 2002].

**Figure 12.**
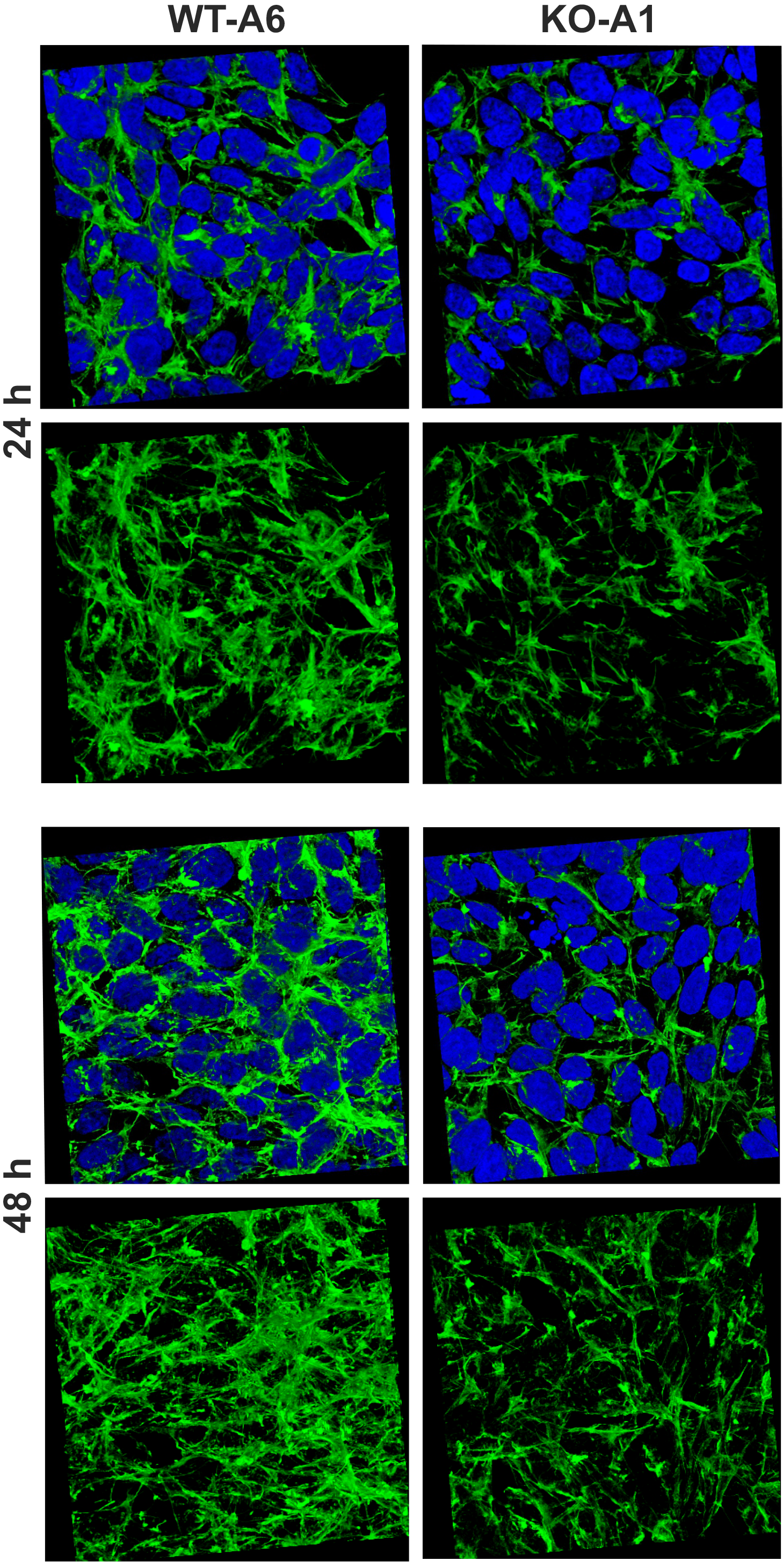
Organization of actin cytoskeleton in the *Setd3*-deficient HAP1 cells visualized by confocal microscopy. Control HAP1 cells (WT-A6) and Setd3-deficinet clonal HAP1 cells (KO-A1) were cultured for either 24 or 48 hours. Localization of filamentous actin was determined by staining with TRITC-phalloidin (green), while nuclei were visualized by Hoechst 33342 (blue). Three-dimensional confocal microscope images of actin cytoskeleton are shown with or without stained nuclei (blue). For 3D analysis, confocal Z-stacks comprising 10-13 optical slices were reconstructed into each 3D image with the aid of Zeiss Zen 2012 software. Representative images are shown, while similar results were obtained with three different HAP1 *Setd3*-deficient clonal cell lines (KO-A1, KO-A3 and KO-A5).

A routine daily inspection of HAP1 cell cultures revealed that at about 80-90% culture confluency, *Setd3*-deficient HAP1 cells tended to acidify IMDM culture medium more quickly than the control ones. This phenotype was rather not attributable to the differences in cell culture density between control and KO cell lines, suggesting an increased formation of lactate by *Setd3*-deficient cells. To test this possibility, both control and *Setd3*-deficient HAP1 cells (90% culture confluency) were incubated in the presence of freshly exchanged IMDM medium for 20h. Spectrophotometric assay was then employed to determine lactate concentration in deproteinized culture medium as described under *“Experimental Procedures”*. As shown in Fig. 13. the formation of lactate in SETD3-KO cell lines was increased by about 15-18% in comparison with that determined in the control cells, indicating a shift of their metabolism towards glycolytic phenotype.

**Figure 13.**
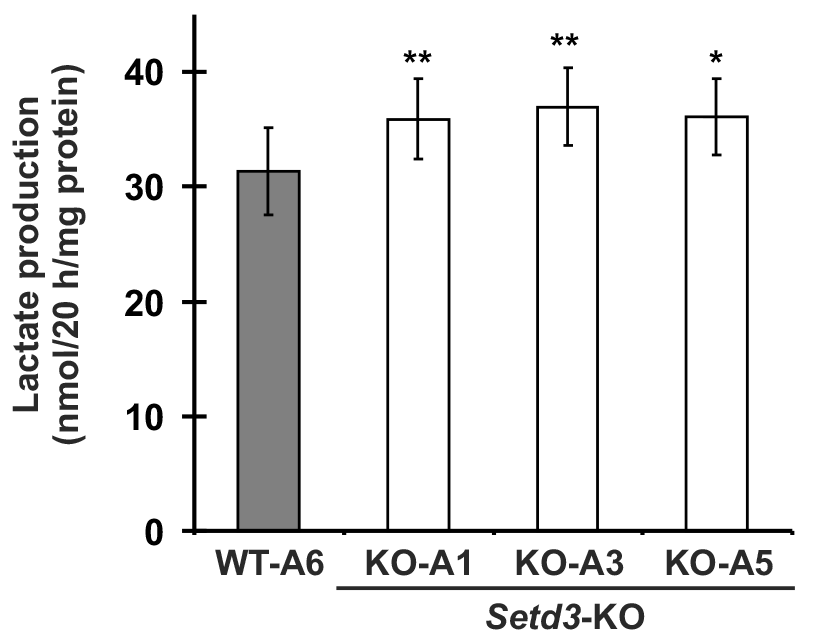
Lactate production in the *Setd3*-deficient HAP1 cells. Experiments were performed employing three HAP1 cell lines deficient in *Setd3* (KO-A1, KO-A3 and KO-A5) and a control HAP1 cell line (WT-A6) that has been submitted to the CRISPR/Cas9 gene inactivation procedure, but yielding no modification of the *Setd3* gene; this was done to exclude off-target effects of the procedure on lactate production. HAP1 cells (≈90% culture confluency) were supplemented with fresh medium and incubated for another 20 hours as described under “Experimental Procedures.” Lactate concentration in samples of deproteinized culture medium was determined spectrophotometrically with the use of lactate dehydrogenase (Noll, 1983). Values are the means ± S.E. (error bars) of 3 independent experiments performed with cells coming form three different culture passages. Statistical significance was analyzed using a paired sample Student’s t-test. *P<0.07, **P<0.01.

## DISCUSSION

### Molecular Identity of Rat Actin-specific Histidine N-Methyltransferase

The presence of two distinct histidine *N*-methyltransferases has been reported in rabbit leg muscle by Raghavan and colleagues [1992]. The first enzyme, carnosine *N*-methyltransferase catalyzes methylation of the histidine imidazole ring of carnosine (β-alanyl-L-histidine), an abundant dipeptide in skeletal muscles of vertebrates. The second one was shown to methylate H73 residue of mammalian β-actin protein. The previous studies performed in our laboratory resulted in the identification of genes coding for carnosine *N*-methyltransferase in vertebrates [Drozak *et al.*, 2013 and 2015], while the current investigation provides data allowing the molecular identification of rat actin-specific histidine *N*-methyltransferase as the SET domain-containing protein 3, disclosing the identity of mammalian actin-methylating enzyme. This conclusion results from the following findings: (i) two different and extensive rounds of purification of actin-methylating activity from rat muscles resulted in the identification of SETD3 protein as the only logical candidate for the enzyme; (ii) the recombinant rat SETD3 catalyzes specific methylation of H73 residue present in both the recombinant β-actin protein and the synthetic peptide corresponding to the nucleotide sensor loop of β-actin; and (iii) in contrast to actin isolated from the control HAP1 cells, the protein present in *Setd3*-deficient cells is essentially devoid of methylated His73 residues, as verified by MS/MS analysis.

Methylation of His73 is highly conserved modification among actins from many different eukaryotes, though it has been found completely absent from actin of *Naegleria gruberi* and *Saccharomyces cerevisiae* [Sussman *et al.*, 1984; Nyman *et al.*, 2002]. The finding that orthologues of SETD3 protein are present in animals, plants and certain fungi, but not in proteomes of the abovementioned proteozoan and yeast species further supports the identification of SETD3 as actin-specific histidine *N*-methyltransferase.

SETD3 was originally reported as methyltransferase catalyzing the modification of histone H3 at Lys4 and Lys36 residues in zebrafish and mouse [Kim *et al.*, 2011; Eom *et al.*, 2011]. Independent studies of Chen and coworkers [2013] confirmed histone methyltransferase activity of SETD3 and suggested that the enzyme may have other non-histone substrates in cytoplasm due the presence of Rubisco LSMT substrate binding domain in its structure. This view was further strengthened by a finding that SETD3 is predominantly localized in the cytosol of human cells, while only small portion of the enzyme is present in the nucleus [Cheng *et al.*, 2017]. Though, we have not examined the activity of recombinant SETD3 towards histones, as isolated histones are extremely poor substrates [Chen *et al.*, 2013], this activity seems to be well-documented and confirmed by several studies. Our present results are therefore complementary to the above-listed findings in that they disclose actin as the cytosolic substrate of SETD3 enzyme (perhaps the most abundant one), and show that SETD3 exhibits plausibly a dual methyltransferase specificity towards both Lys and His residues of proteins.

### Quasi-native β-actin from Bacterial Inclusion Bodies is a Good Substrate for Actin-specific Histidine N-methyltransferase

The molecular identification of the actin-specific histidine *N*-methyltransferase thorough its extensive purification from rat muscles was only possible, owing to the fact that we have had large amounts of pure β-actin monomers for radiochemical enzymatic assays. Though β-actin was produced and subsequently purified from inclusion bodies of *E. coli* in the nucleotide-free form, it acted as an efficient substrate for the methyltransferase studied. In view of the facts that (i) it is impossible to produce natively folded actin in bacteria [Stemp *et al.*, 2005; Martín-Benito *et al.*, 2002] and (ii) ATP/ADP-free actin denatures rapidly and irreversibly [Graceffa and Dominguez, 2003], it seems that actin preparation used in the present study may comprise a sort of actin molecules, exhibiting stable non-native spatial structures. Indeed, the actin methylation reaction was still substoichiometric, and we have observed methylation of up to 35% of the total actin present in the reaction mixture. This effect was readily reproducible in the presence of bacterial SAH nucleosidase and adenine deaminase, indicating that it was not due to the potential accumulation of SAH in the reaction mixture (unpublished data, Drozak *et al.*). This indicates that only small portion of actin monomers produced in bacteria and purified under denaturing conditions have conformation that could be recognized by the SETD3 enzyme. This statement is in line with a previous report, showing only minor methylation of the inclusion body actin (up to 1% protein) by rabbit actin methyltransferase [Raghavan *et al.*, 1992]. Furthermore, SETD3 was at least 500-fold less active towards the synthetic peptide corresponding to the amino acid sequence surrounding the H73 residue in comparison with the recombinant β-actin, indicating that other structural features of the actin molecule than the primary sequence of the sensor loop are likely important for its recognition by the methyltransferase.

Both tested SETD3 orthologs are rather sluggish enzymes, with a *k*_cat_ of 0.6-0.8 × min^-1^, resembling values estimated for protein methyltransferases rather than that measured for the enzymes catalyzing methylation of low molecular weight metabolites (Clarke and Banfield, 2001, Osborne *et al.*., 2007). Considering that the intracellular concentrations of SAM and β-actin monomers in vertebrate cells are about 30 μM and 50-100 μM, respectively [Clarke and Banfield, 2001; Pollard, 2017], both rat and human SETD3 exhibited a high affinity for both SAM (K_M_ ≈ 0.3 μM) and β-actin (K_M_ ≈ 0.8 - 3 μM). These observations suggest that the enzymes may be fully saturated by substrates and operate at rates that are close to their V_max_ *in vivo*. However, it should be pointed out that the recombinant β-actin used in the present kinetic studies is certainly not a physiological form of this substrate. Therefore, the affinity of enzyme to β-actin remains to be verified.

It is very intriguing, that in our hands both the recombinant ATP- and ADP-β-actins produced in yeast (*S. cerevisiae*), which are expected to have a native conformation, were not methylated by the SETD3 enzyme. This may explain the fact, that the molecular identity of actin-methylating enzyme has remained unidentified so far. On the other hand, more interestingly, this finding may suggest that the presence of ATP or ADP could create a structural hindrance for SETD3 to reach H73 residue, if the methyltransferase interacts with actin through the nucleotide-binding cleft (cf. Fig. 6). Therefore, we have prepared a stable recombinant nucleotide-free β-actin in complexes with either profilin or cofilin and found that only very low amounts of the cofilin-β-actin (up to 3-4% of total protein) were methylated by the SETD3 enzyme. For unknown reasons, however, the extent of methylation was barely reproducible among different preparations of these protein complexes (unpublished data, Drozak *et al.*). Taken together, these observations have led us to hypothesize that the nucleotide-free actin monomer forming complexes with an actin-binding protein of unknown yet identity might be the “true” substrate for the SETD3 methyltransferase. If correct, it resembles the activity of yeast YIL110W protein, the only other known protein histidine *N*-methyltransferase up to date that methylates H243 residue of ribosomal Rpl3 protein under conditions when this protein is associated with the ribosome [Webb *et al.*, 2010; Al-Hadid *et al.*, 2014].

### Biological Importance of SETD3 Methyltransferase

Several reports describing the role of SETD3 in the regulation of cell cycle and apoptosis [Kim *et al.*, 2011], myocytes differentiation [Eom *et al.*, 2011], cell response to hypoxic conditions [Cohn *et al.*, 2016] and tumorigenesis [Pires-Luis *et al.*, 2015; Cheng *et al.*, 2017] have been published so far. Some of these studies identified methylation of histones as a primary mechanism of SETD3 action [Kim *et al.*, 2011; Eom *et al.*, 2011], while others suggested that the enzyme may work rather by modifing nonhistone substrates present in the cytosol [Chen *et al.*, 2013, Cohn *et al.*, 2016]. The existence of two distinct domains in the enzyme, i.e. SET and Rubisco LSMT substrate binding, has been suggested to be the structural basis for such dual substrate specificity [Chen *et al.*, 2013].

Our results presented here identify β-actin as a novel substrate of SETD3 and show that disruption of its activity leads to the depletion of F-actin and a loss of cytoskeleton integrity. This phenotype is most likely due to an increase in the rate of depolymerization of actin filaments, as it was previously shown for a hypomethylated recombinant actin *in vitro* [Nyman *et al.*, 2002]. The inherited instability of F-actin in *Setd3*-defficeint cells is plausibly also the primary cause for an increased lactate production by these cells. Both polymerization and depolymerization of actin filaments occur continuously even in resting cells and constitute a major energy drain, causing up to 50% of total ATP consumption in vertebrate cells [Kudryashov and Reisler, 2012]. Thus, accelerated degradation of F-actin is likely to drive increasing demand for ATP that could be satisfied by shifting the cell metabolism towards glycolytic one. The phenotype of *Setd3*-defficeint cells is therefore somewhat similar to that of malignant cells, suggesting that the SETD3 activity may play a role in suppressing tumor development. Indeed, hypomethylation of actin was observed in rat cells following their transformation with the Src oncogen [Chiou et al., 2012]. Furthermore, enzymatically inactive SET domain-loss SETD3 was shown to function as a dominant negative mutant, promoting oncogenesis in human cells [Chen *et al.*, 2013], most likely by blocking the methylation of unidentified SETD3 substrate. It is tempting to speculate that actin protein might be this substrate.

Taken together, we suggest that SETD3 exerts pleyotropic activity, regulating many different and not necessary related processes, depending on the site of its action, i.e. cytosol or nucleus. This notion is supported to some extent by following findings: (i) SETD3 was shown physically to interact with about 170 different proteins and at least some of them are plausibly its substrates [Cohn *et al.*, 2016] and (ii) a loss of SETD3 activity induces rather complex pattern of disturbances in a mammalian organism, as reported for *setd3*-deficient mice that suffer from multisystemic complications, affecting nervous, cardiovscular, endocrine, reproductive and skeletal systems as well as the metabolism (Mouse Genome Informatics database, MGI:1289184).

### Conclusions

In the current investigation, we have identified the rat actin-specific histidine *N*-methyltransferase as SETD3 protein, an enzyme catalyzing the methylation of H73 residue in the β-actin protein. The SETD3 enzyme is therefore the first protein-histidine methyltransferase identified in vertebrates so far and the first SET-domain containing enzyme, exhibiting a dual methyltransferase specificity towards both Lys and His residues of proteins. As the loss of SETD3 in HAP1 cells induces phenotypic changes resembling those present in cancer cells, we hypothesize that hypomethylation of actin might be involved in tumorigenesis. Finally, this work also shows that non-native protein substrates might be useful tools in search for novel protein methyltransferases.

## ACKNOWLEDGMENTS

This investigation was partially financed by the Opus-6 grant (DEC-2013/11/B/NZ1/00078) from the Polish National Science Centre and DSM 501-D114-86-011500-23 from the Polish Ministry of Science and Higher Education. M.T. was an Erasmus+ trainee supported by the European Union. Research was carried out with the use of CePT infrastructure financed by the European Union – the European Regional Development Fund within the Operational Program “Innovative economy” for 2007-2013. Work in the laboratories of M.V.D.C. and D.V. was supported by a grant from the Fonds National de la Recherche Scientifique (FNRS, Belgium).

The authors would like to express their appreciation to Professor Jadwiga Bryla (University of Warsaw), who read the manuscript carefully and provided numerous helpful suggestions.

## ABBREVIATIONS

The abbreviations used are: SETD3; SET domain-containing protein 3; MS/MS: tandem mass spectrometry; SAH: *S*-adenosyl-L-homocysteine; SAM: *S*-adenosyl-L-methionine.

## AUTHOR CONTRIBUTIONS

J.D. conceived and designed the study and wrote the paper.

J.D., S.K, A.K.S, M.T., T.I. and A.K.J. performed and analyzed experiments shown in Tables 1-3, 5, and Figures 1-10 and 13.

M.V.C., D.V. and A.K.S designed, performed and analyzed experiments, aiming at the generation of *Setd3*-deficient cells and MS/MS analysis of intercellular actin (Figure 11, Table 4).

IG designed, performed and analyzed experiments shown in Figure 12.

All authors reviewed the results and approved the final version of the manuscript.

